# Activity-based chemical proteomics uncovers unexpected covalent targets of E64d and reveals a role for cysteine cathepsins in PLD3 proteostasis

**DOI:** 10.64898/2026.08.14.744826

**Authors:** Manuel Hertwig, Pavel Kielkowski

## Abstract

Catalytic activity of 5′-3′ exonuclease Phospholipase D3 (PLD3) is associated with immune signaling and neurodegeneration including Alzheimer’s disease. PLD3 undergoes multiple post-translational modifications and proteolytic cleavage to establish its catalytically active form. However, the proteases catalyzing the cleavage of PLD3 have remained unidentified. To study the proteolytic cleavage of PLD3, we have evaluated the small molecule covalent inhibitor E64d that blocks proteolysis catalyzed by cysteine cathepsins. To validate the selectivity of E64d, we have designed and synthetized an E64d propargyl analogue and carried out a detailed activity-based protein profiling to reveal a broad engagement of the compound with other protein targets including bleomycin hydrolase (BLMH), Kelch-like ECH-associated protein 1 (KEAP1), transcription elongation factor SPT5 (SUPT5H) and asparagine synthetase (ASNS). The specificity of the E64d-protein interactions was confirmed by biochemical assays and mass spectrometry-based site identifications. In neurons, treatment with E64d lead to about 50-fold PLD3 accumulation and dysregulation of its proteolytic cleavage, while there was only a minor overall change on the whole proteome level. Taken together, this study provides insights into previously unknown E64d selectivity and renders cysteine cathepsins responsible for PLD3 degradation in neurons. It highlights the importance of cysteine cathepsins activity in neuronal lysosomes for proper PLD3 processing and hence it suggests that their activation might be responsible for decreased PLD3 levels in neurons of patients with Alzheimer’s diseases. These findings are key for further elucidation of PLD3 function in neurodegenerative diseases.

## Introduction

Proteolytic cleavage is a fundamental post-translational modification that regulates protein function by irreversibly altering protein structure, activity, localization, and stability. It is essential for processes such as signal transduction, apoptosis, and protein maturation, thereby maintaining protein homeostasis^[1]^. Disruption of protein homeostasis is a hallmark of neurodegenerative diseases, leading to the accumulation of misfolded or aggregated proteins that impair neuronal function^[2–5]^. Cysteine cathepsins, a family of lysosomal proteases, play a critical role in maintaining proteostasis by mediating protein degradation and turnover within the endolysosomal pathway^[6]^. Dysregulation of cathepsin protease activity has been implicated in disorders such as Alzheimer’s disease and Parkinson’s disease, highlighting their potential as therapeutic targets in neurodegeneration^[7,8]^. Among cathepsin inhibitors, E64d is a well-established broad-spectrum cysteine protease inhibitor due to its ability to penetrate cells and reduce catalytic activity of cysteine cathepsins by covalently binding their catalytic cysteine in the active site (**Figure 1a** and **b**)^[9–12]^. Because of its efficient inhibition of cysteine cathepsins, E64d has been tested for treatment of various diseases including muscular dystrophy^[13]^, heart failure^[14]^, SARS-CoV-2^[15]^ and Alzheimer’s diseases^[7]^. However, its clinical translation has been limited by suboptimal pharmacokinetic properties, concerns regarding on-target effects associated with long-term protease inhibition, and a lack of understanding regarding its target selectivity.

**Figure 1.**
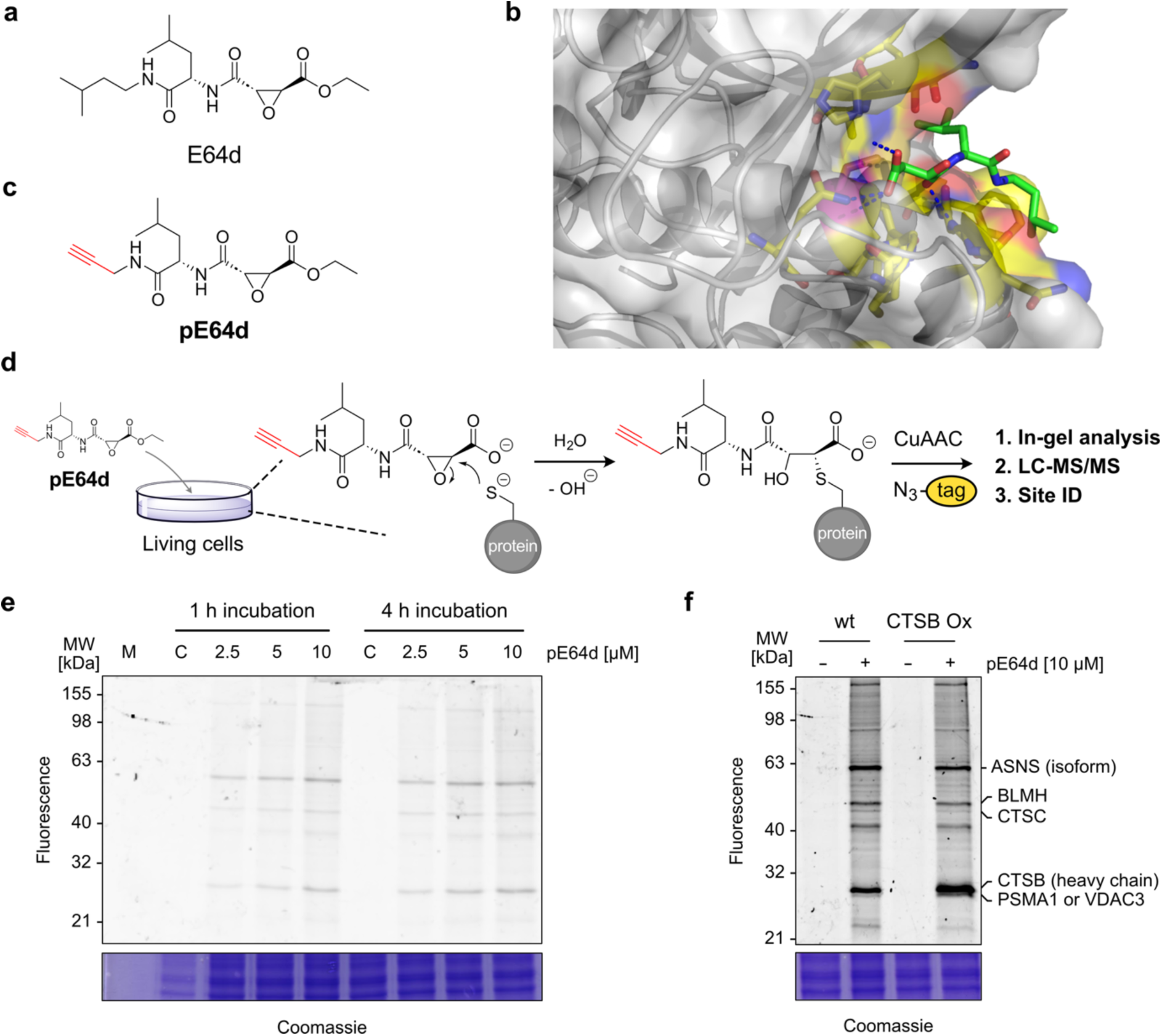
Design of E64d chemical proteomics probe. **a**) E64d structure. **b**) Crystal structure of CTSB active site with E64c. The catalytic cysteine is highlighted in pink. PDB 1ITO^[29]^. **c**) Structure of the chemical proteomics probe pE64d. **d**) E64d mechanism of action leading to a covalent bond between pE64d and reactive cysteines of proteins including cysteine cathepsins. **e**) In-gel fluorescence analysis of pE64d protein labelling in HEK293T cells after indicated treatment times and concentrations. C - DMSO control. **f**) In-gel fluorescence analysis of protein labelling by pE64d in HEK293T cells without and with transiently overexpressed CTSB. The putative protein targets were assigned based on the molecular weights of proteins identified after enrichment and LC-MS/MS analysis. CuAAC: Cu(I)-catalyzed azide-alkyne cycloaddition.

Interestingly, cysteine cathepsins have been considered to be involved in the degradation of the N-terminal transmembrane domain of PLD3 after its proteolytic processing ^[16]^. PLD3 has emerged as a promising therapeutic target in Alzheimer’s disease^[17–20]^. PLD3 is a lysosome-associated transmembrane protein highly expressed in neurons and is thought to contribute to endolysosomal homeostasis through nucleic acid metabolism via its reported 5′-3′ exonuclease activity^[21–23]^. Genetic and functional studies have linked PLD3 dysfunction to Alzheimer’s disease through decreased expression levels and rare variants that have been associated with increased amyloid pathology and impaired neuronal function^[17,24–26]^. PLD3 is synthesized in the endoplasmic reticulum and trafficked through the Golgi apparatus to endolysosomal compartments, where it undergoes extensive post-translational modifications^[16,27]^. Proteolytic cleavage generates a mature lysosomal (soluble) form of PLD3, which is necessary to form the catalytically active homodimer^[22,28]^. In addition, PLD3 undergoes post-translational modifications including *N*-linked glycosylation and AMPylation, a reversible modification that regulates proteolytic processing^[27,28]^. While healthy neurons display the highest cellular levels of soluble PLD3 in comparison to other cell types in which PLD3 is present as the full-length protein in significantly lower levels, dysregulation of PLD3 processing was recently observed in Parkinson’s patient-derived neurons^[28]^. Despite increasing interest in PLD3, the molecular mechanisms governing its processing, trafficking, and regulation remain poorly defined. The initial report has suggested that CTSB might only digest the N-terminal peptide, but it is not responsible for the cleavage that releases the soluble part of PLD3^[16]^.

Here, we set out to utilize activity-based protein profiling (ABPP) to evaluate the selectivity of E64d and to assess the impact of cysteine cathepsin inhibition on PLD3 processing in HEK293T cells and iPSC-derived neurons during neuronal differentiation.

## Results

Given the therapeutic potential of E64d, we set out to evaluate the selectivity of this covalent inhibitor and to identify its putative off targets. In parallel, using an activity-based E64d probe, we sought to identify proteases outside of the cysteine cathepsin family that might be covalently trapped by E64d and therefore could be responsible for PLD3 cleavage. Previously, E64d was shown to inhibit PLD3 processing, but validation of the results in knockout cell lines of cysteine cathepsins provided rather unclear outcomes^[16]^. Based on the crystal structure of the CTSB active site with E64c, the free acid analogue of E64d (**Figure 1b** and Figure S1), we have designed the propargyl containing analogue pE64d (**Figure 1c**)^[29]^. This allows to covalently trap reactive cysteines in protease active sites and provides a terminal alkyne for downstream analysis by in-gel fluorescence or affinity purification followed by LC-MS/MS proteomics (**Figure 1d**). The three-step synthesis (Figure S2) has provided the pE64d probe, which was then used for treatment of HEK293T cells. For the in-gel fluorescence analysis, the pE64d probe labelled proteins were conjugated with a TAMRA-azide using the Cu(I)-catalyzed azide-alkyne cycloaddition (CuAAC) under strongly reducing conditions to avoid any chemical side-reactivity^[30–32]^.

The optimization of the probe concentration and treatment time was done using the in-gel fluorescence analysis to show the first protein labelling at 2.5 μM of pE64d final concentration in cell culture medium and with increasing fluorescence intensity towards 10 μM (**Figure 1e**). The appearance of the fluorescence bands at around 30 and 50 kDa suggested the labelling of some of the cysteine cathepsins, which have a comparable molecular weight (CTSB ∼30 kDa and CTSC ∼50 kDa). Further increase of pE64d concentration (up to 500 μM) led to stronger protein labelling with the same pattern (Figure S3). To analyze dynamics of pE64d labelling, pulse-chase labelling was performed using 10 μM pE64d to show relatively stable protein labelling for about 18 h after probe-free media exchange, and nearly complete disappearance of the labelling after three days (Figure S4). In parallel to the pE64d testing in HEK293T, the probe labelling was examined in near-haploid cells derived from patient with chronic myeloid leukemia (Hap-1) showing the same labelling pattern (Figure S5). With the focus on CTSB catalytic activity and known probe reactivity, the HEK293T cells were transiently transfected with a CSTB gene containing plasmid before pE64d treatment. The subsequent in-gel fluorescence analysis revealed a strong increase of fluorescence at around 30 kDa corresponding to the active CTSB (**Figure 1f**). Next, to identify putative off-targets of E64d, we have proceeded with pull-down of pE64d labelled proteins using a biotin-azide tag and SP2E workflow, which is composed of two steps utilizing carboxylate modified magnetic beads for protein clean-up and then streptavidin-coated magnetic beads for affinity-based enrichment of probe labelled proteins^[33]^. The SP2E pull-down of labelled proteins was done from four different HEK293T cell cultures, treated with 10, 50 or 100 μM pE64d probe. The subsequent calculation of the proteomics data by DIA-NN^[34]^ and differential expression analysis with limma and limpa^[35,36]^ showed a consistent enrichment of CTSB across the used pE64d probe concentrations (**Figure 2a-c**). Interestingly, 10μM pE64d treatment not only led to significant enrichment of cytosolic cysteine protease BLMH, but also proteins of diverse molecular functions including a transcription factor (SUPT5H), a ligase (ASNS), as well as scaffolding and adapter proteins (KEAP1 and PSMA1) (**Figure 2a** and Figure S6). Of note, KEAP1 contains a highly reactive cysteine residue reported by Bar-Peled *et al.*^[37]^. Treatment with 50 µM and 100 µM resulted also in significant enrichment of Interleukin-1 receptor-associated kinase 1 (IRAK1), which is a crucial kinase of the toll-like receptor signaling pathway. Overall, a group of six proteins was consistently enriched across all concentrations (**Figure 2d**). A more detailed analysis of enriched proteins showed also a highly consistent enrichment over the four replicates (**Figure 2e**), which is supported by profile plot analysis of the hit proteins (**Figure 2f**). Taken together, the in-gel fluorescence analysis and LC-MS/MS proteomics of pE64d probe enriched proteins confirms the engagement of E64d with CTSB, but also uncovers a group of previously unknown proteins, that directly interact with the E64d scaffold. The engagement with other proteins than cysteine cathepsins might provide the cues for E64d side-effects.

**Figure 2.**
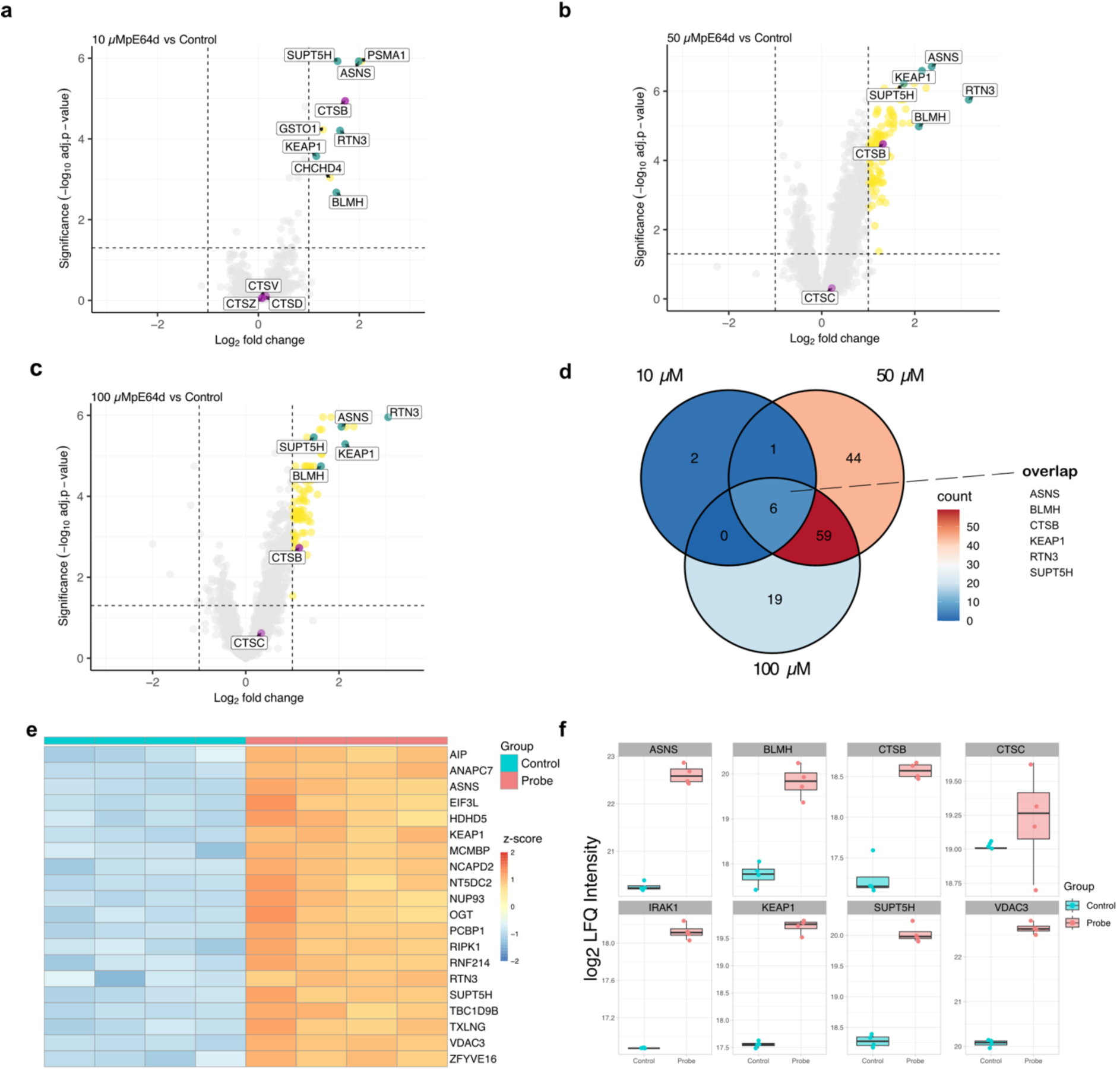
Mass spectrometry-based proteomics analysis of pE64d targets in HEK293T cells. **a-c**) Volcano plots visualizing the concentration dependent pE64d protein enrichment. Significantly enriched proteins are represented by yellow circles (*n* = 4 biological replicates, cut-off p-value < 0.05, and fold-change > 1). Cysteine cathepsins are depicted in magenta. The group of the most significantly enriched proteins across various conditions is depicted in green. **d**) Venn diagram showing the overlap of significantly enriched proteins at different concentrations. **e**) Heatmap summarizing the top 20 most significantly enriched proteins by adj.p-value using 50 μM pE64d. **f**) Profile plots of selected significantly enriched proteins after treatment with 50 μM pE64d (*n* = 4 biological replicates).

To validate the enrichment of the proteins using a mass spectrometry-independent method, we have selected CTSB and a group of another four proteins including BLMH, PSMA1, SUPT5H and IRAK1 for pull-down of pE64d labelled proteins from cell lysates. Instead of their tryptic on-beads digest, the proteins were eluted from the beads, separated on sodium dodecyl sulfate-polyacrylamide gel electrophoresis (SDS-PAGE) and visualized by western blot using corresponding antibodies. Of note, CTSB was overexpressed in HEK293T cells because endogenous levels were not detectable with the corresponding antibody. All selected proteins showed a clear probe dependent enrichment confirming their direct interaction and reactivity with the pE64d probe (**Figure 3a** and **b** and Figure S7).

**Figure 3.**
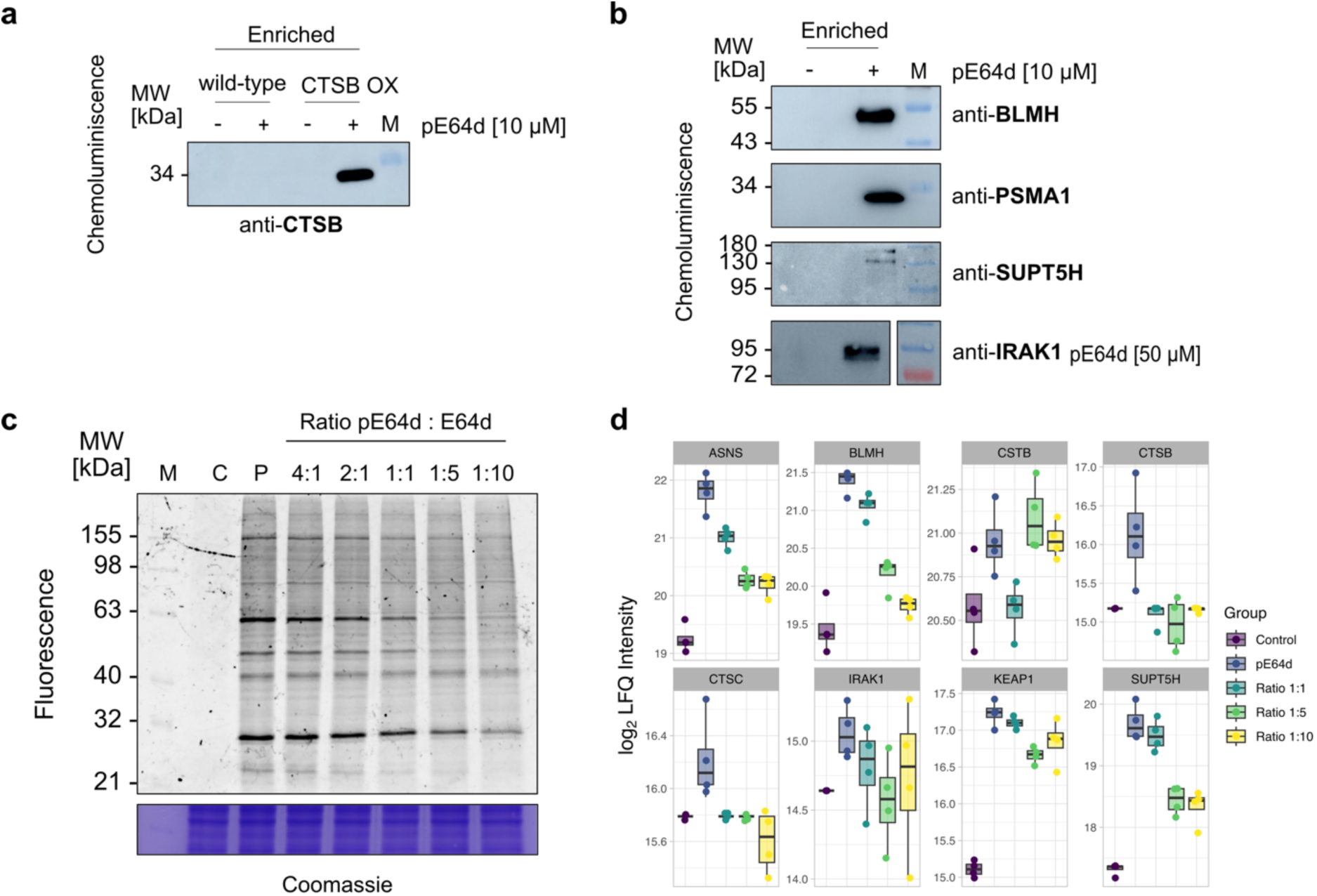
Validation of pE64d targets. **a**) Western blot of the pE64d enriched proteins from HEK293T cells without or with CTSB transient overexpression. **b**) Western blot using various antibodies of the pE64d (10 µM or 50 µM) enriched proteins from HEK293T cells. **c**) In-gel fluorescence analysis of the competition experiment between pE64d probe and the parent compound in HEK293T cells. C - DMSO control, P - pE64d 10 µM. **d**) Profile plots visualizing the MS-based proteomic results after enrichment with pE64d and competition with the parent compound (*n* = 4 biological replicates).

Next, to examine whether the enriched proteins engage with the pE64d probe because of E64d scaffold recognition or because of the epoxide reactivity with nucleophilic cysteine residues, we have carried out a competition experiment between the pE64d probe and the parent compound E64d in HEK293T cells, in which the concentration of pE64d probe was kept constant and competed with increasing concentrations of E64d. The resulting protein labelling was analyzed by in-gel fluorescence and after pull-down by LC-MS/MS proteomics. The in-gel analysis revealed that most of the major fluorescence bands decrease in intensity with increasing addition of parent compound E64d, but not all (**Figure 3c**). The quantitative mass spectrometry-based experiment then led to deconvolution of dose-dependent responsive targets that included CTSB, ASNS, BLMH, CTSC and SUPT5H, but not cystatin-B (CSTB), an endogenous CTSB inhibitor, KEAP1 and IRAK1 (**Figure 3d**). Together, these experiments validate the fidelity of mass spectrometry-based chemical proteomic experiments and off-target reactivity of E64d.

Although the reactivity of E64d is well-established for cysteine cathepsins, the modification sites on the newly identified target proteins were unknown. Therefore, we proceeded with a pull-down of pE64d-labelled proteins in wild-type or CTSB overexpressing HEK293T cells using a desthiobiotin-azide linker, which facilitates the elution of pE64d-modified peptides from streptavidin-coated magnetic beads due to its weaker binding affinity compared to biotin (**Figure 4a**). The modified peptides were expected to appear with a mass shift of 696.38 Da (**Figure 4b**). High resolution MS^2^ spectra were acquired using data-dependent acquisition and the obtained spectra were searched in MSFragger with open, closed and offset search modes. This confirmed labelling of CTSB and CTSC catalytical cysteines (**Figure 4c, d**), and identified the binding sites on SUPT5H and PSMA1 (**Figure 4e** and **Figure S8**). A complementary search for diagnostic ions, including the desthiobiotin-oxonium ion and probe-specific cysteine-pE64d fragments, further corroborated the fidelity of searches (**Figure 4b**, Figure S9 and Tables S1 and S2). Importantly, no modified peptides were found in the control group that was not treated with the pE64d probe (Figure S10). In total 144 sites were identified with 19 of them overlapping with protein-level enrichment using 50 μM pE64d (**Figure 4f**). Because the modification site on IRAK1 was not initially found, we conducted an IRAK1 antibody-based immunoprecipitation experiment, which lead to desired site identification and the gave a mass shifts of 282.1215 for pE64d on cysteine 578 **(Figure S8**). To further corroborate the fidelity of the probe and reactivity of CTSB catalytic cysteine, HEK293T cells transiently overexpressing wild-type CTSB or CTSB C108A mutant were treated with pE64d and enriched on streptavidin beads. The western blot analysis indeed confirmed the selectivity of the probe towards the catalytic cysteine C108, while no enrichment of CTSB could be observed in the CTSB C108A overexpressing cells (**Figure 4g** and **Figure S11**). Interestingly, pE64d reacts with several previously described reactive cysteines including C578 on IRAK1 (in isoform 2, it corresponds to C610 of the canonical sequence, **Figure S8**) and C622 on SUPT5H^[38]^. Together, the site-ID experiments validated the pE64d off-targets identified by initial pull-down experiments.

**Figure 4.**
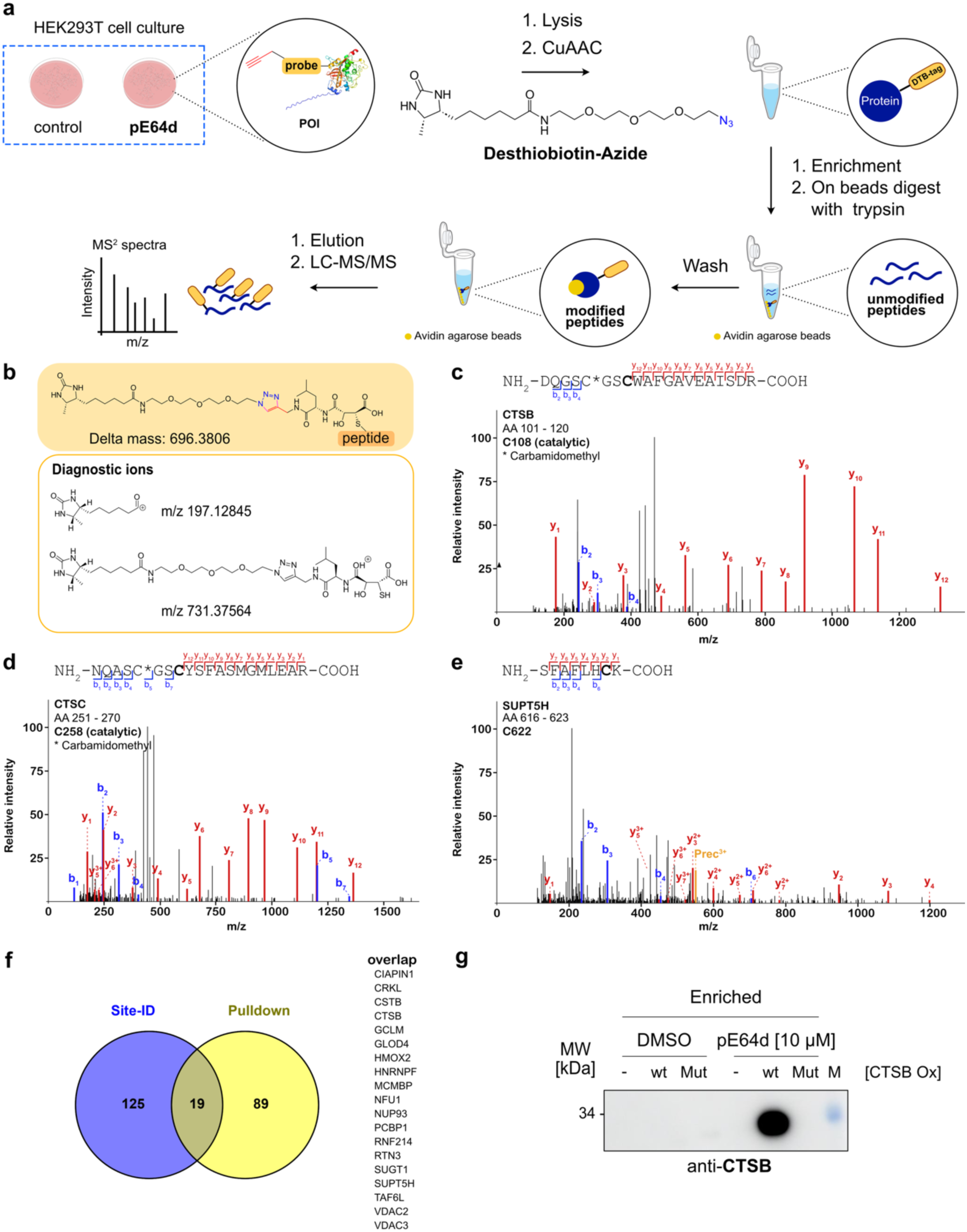
Identification of pE64d binding sites. **a**) Scheme showing the site identification workflow and structure of desthiobiotin-azide linker. **b**) Structure of the pE64d probe modified peptides with the delta mass used for their search. **c**-**e**) Examples of MS2 spectra of pE64d modified cysteines found by FragPipe. **g**) Overlap between significantly enriched proteins shown in Figure 2b using 50 μM pE64d probe and site-ID experiment using the same probe concentration. **f**) Western blot using anti-CTSB antibody against the pE64d enriched proteins from HEK293T cells without or with wild-type or C108A mutant CTSB transient overexpression.

Having characterized the off-target landscape of E64d in HEK293T cells and found no other lysosomal protein proteases other than cysteine cathepsins, we turned our focus to the analysis of E64d activity during neuronal differentiation and its impact on PLD3 processing. The pE64d probe was utilized in human induced pluripotent stem cells (hiPSCs), which feature a doxycycline-inducible expression of Neurogenin-1 and Neurogenin-2, enabling rapid and robust differentiation into a homogeneous culture of dopaminergic neurons (iNGNs)^[23,39]^. The activity of E64d was probed at five different time points during differentiation including the initial hiPSCs and ten-day neurons (**Figure 5a**). To confirm successful differentiation of hiPSCs into neurons we measured the whole proteome of stem cells and nine-day old neurons. We confirm differential protein expression, appearance of neuronal markers and decrease of stem cell markers. Gene ontology enrichment and gene set enrichment analysis (GSEA) supported the neuronal cell state (**Figure S12**). We then went on to investigate expression levels and enrichment of individual proteins. First, we detected an increase of CTSB levels during neuronal differentiation by western blot, while CTSZ levels steadily decreased (**Figure 5b**)^[40]^.

**Figure 5.**
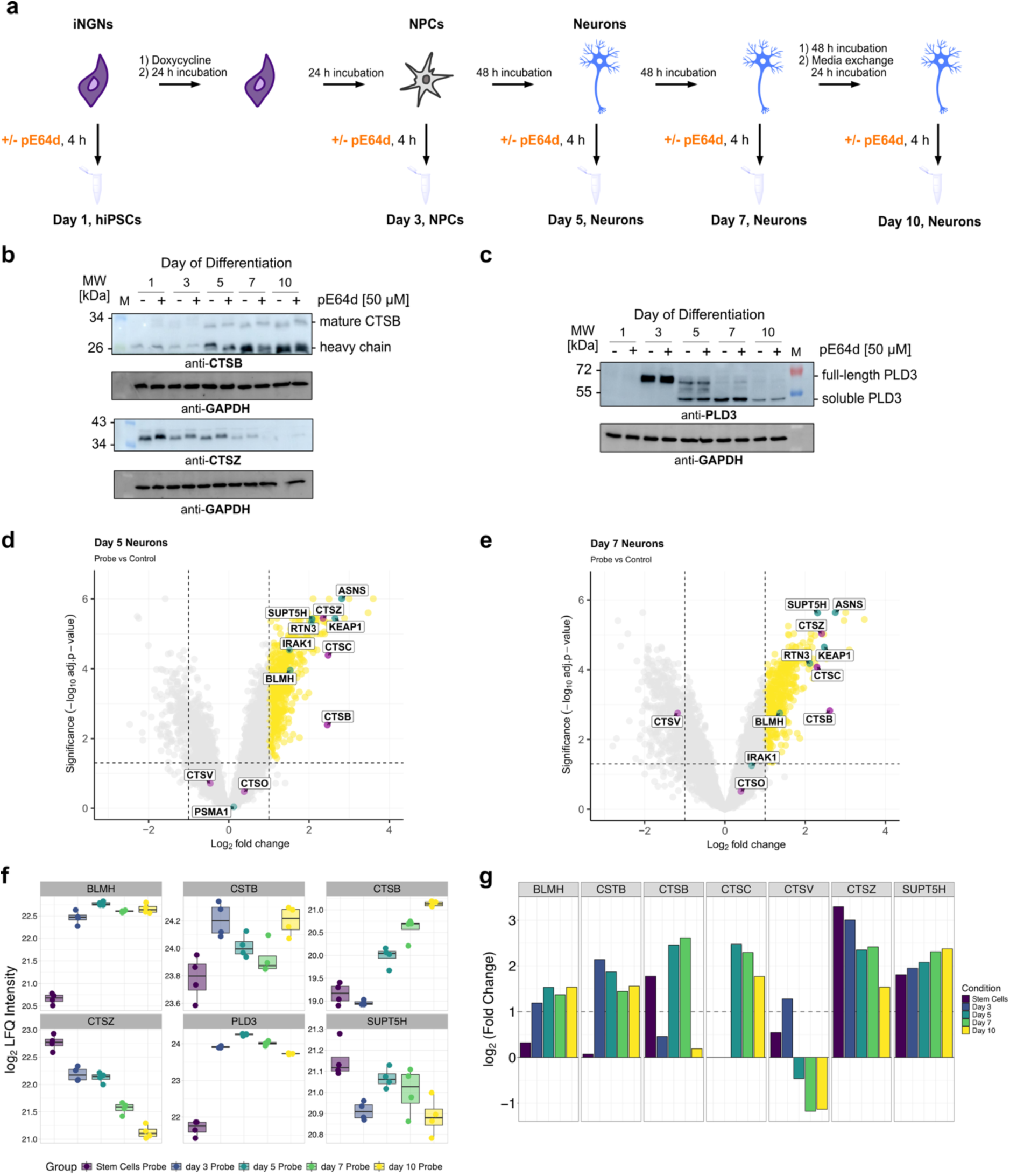
Profiling of pE64d protein targets during iNGNs neuronal differentiation. **a**) Scheme showing the iNGN differentiation timeline, pE64d treatments and sample collection. **b**) Western blot using anti-CTSB and anti-CTSZ antibodies to characterize their levels during neuronal differentiation. **c**) Western blot using anti-PLD3 antibody to characterize PLD3 levels and processing during neuronal differentiation. **d**-**e**) Volcano plots showing the pE64d (50 μM) enriched proteins at day 5 and 7 of neuronal differentiation. Significantly enriched proteins are represented by yellow circles (*n* = 4 technical replicates, cut-off p-value < 0.05, and fold-change > 1). Cysteine cathepsins are depicted in magenta. The group of the most significantly enriched proteins across various conditions is depicted in green. **f**) Profile plots showing the expression changes of selected proteins (*n* = 4 technical replicates) from whole cell lysates. **g**) Bar plots visualizing the enrichment ratio (log_2_ (Fold Change) for selected proteins during neuronal differentiation.

Second, we have shown that PLD3 increases rapidly during neuronal differentiation and is progressively proteolytically cleaved into its soluble form, requiring increased proteolytic activity (**Figure 5c**)^[23,28]^. Therefore, we anticipated that treatment with pE64d probe would lead to decreased proteolytic processing of PLD3 if some of the cysteine cathepsins are involved in its cleavage. However, there was no observable change in the amount of the soluble forms between pE64d treated and control cells after 4 h treatment (**Figure 5c**). Third, the pull-down experiment of pE64d labelled proteins during neuronal differentiation exhibited overlapping labeling with HEK293T cells by significant enrichment of ASNS, KEAP1, SUPT5H, BLMH and RTN3 (**Figure 5d-e**). Whole proteome analysis of differentiating neurons by LC-MS/MS confirmed steadily increasing levels of CTSB and continuously decreasing levels of CTSZ, which is in line with the western blot results (**Figure 5f**). Moreover, the whole proteome analysis revealed that treatment with 50 µM pE64d for 4 h did not alter the overall protein levels of investigated proteins and it confirmed successful neuronal differentiation (Figure S13). Interestingly, CTSB was significantly enriched in iNGN stem cells, day 5 and 7, but not in day 3 and 10 neurons suggesting dynamic changes in regulation of the CTSB catalytic activity during differentiation. In contrast, CTSZ was consistently enriched with a decreasing tendency over the differentiation course, following its expression levels (**Figure 5f-g** and Figure S14-S15). CTSC was enriched at day 5, 7 and 10 (**Figure 5g**). Similar to HEK293T cells, no other lysosomal proteases were identified. The PCA of enriched proteins showed clustering of replicates within each group, demonstrating the robustness of the data (Figure S16). Together, the pE64d probe allowed us to profile cysteine cathepsin catalytic activity during neuronal differentiation and hence to directly correlate their abundance with activity to provide important experimental evidence about these essential lysosomal resident proteins^[2,41,42]^. In particular, the drop of CTSZ levels and activity during differentiation, which is complemented by lack of CTSB activity in 10-day iNGN neurons, renders CTSC as the main active cysteine cathepsin protease in mature iNGN derived neurons.

As the four hours treatment with pE64d of neurons showed no inhibition of PLD3 processing, we were wondering if a prolonged treatment with E64d or pE64d might still be efficient in the inhibition. Therefore, we have selected day 5 neurons, which are characterized by strongest transition between full-length and soluble PLD3 suggesting a reprogramming of the PLD3 function, for a treatment with E64d and pE64d. Neurons were treated on day 4 for 30 h with E64d or pE64d, and PLD3 cleavage was analyzed by antibodies recognizing either only the full-length protein (N-terminal antibody) or both the full-length and the soluble protein (luminal antibody). The treatment with E64d or pE64d for 30 h showed a strong increase of full-length glycosylated PLD3 stained by N-terminal antibody (**Figure 6a**). The complementary staining by antibody recognizing the luminal part of PLD3 showed an overall increase of the soluble unmodified and the full-length unmodified PLD3, whereas the soluble unmodified form was increased by two-fold after E64d addition and ∼ 2.5-fold after treatment with pE64d, respectively (**Figure 6b** and Figure S17). Given this large accumulation of different PLD3 forms in day 5 iNGN neurons, we were wondering if these intermediate forms are glycosylated. Therefore, the day 5 iNGN lysates were treated with PNGase F to cleave the N-glycans. Resulting western blots using the N-terminal antibody confirmed more than 20-fold increase of full-length protein (**Figure 6c**), while the proteolytically cleaved soluble PLD3 was increased by ∼ 50-fold (**Figure 6d**). These results suggest that the processing of full-length PLD3 to the cleaved soluble form is not catalyzed by cysteine cathepsins, namely CTSB, CTSC and CTSZ, in 5-day iNGNs. However, the blockage of these cathepsins leads to PLD3 accumulation. In order to determine whether E64d treatment leads to a global change in the proteome by reducing the proteolytic capacity of lysosomes, we have carried out an LC-MS/MS whole proteome analysis from the same control, E64d and pE64d treated 5-day iNGN neurons. Interestingly, we observed only about two-fold increase in PLD3 upon the E64d treatment, which might be accounted to abundant N-glycosylation of the protein in E64d and pE64d treated neurons, which changes the retention times, ionization efficiency and mass to charge ratio of the PLD3 tryptic peptides and hence hampers the correct quantification (**Figure 6e**).

**Figure 6.**
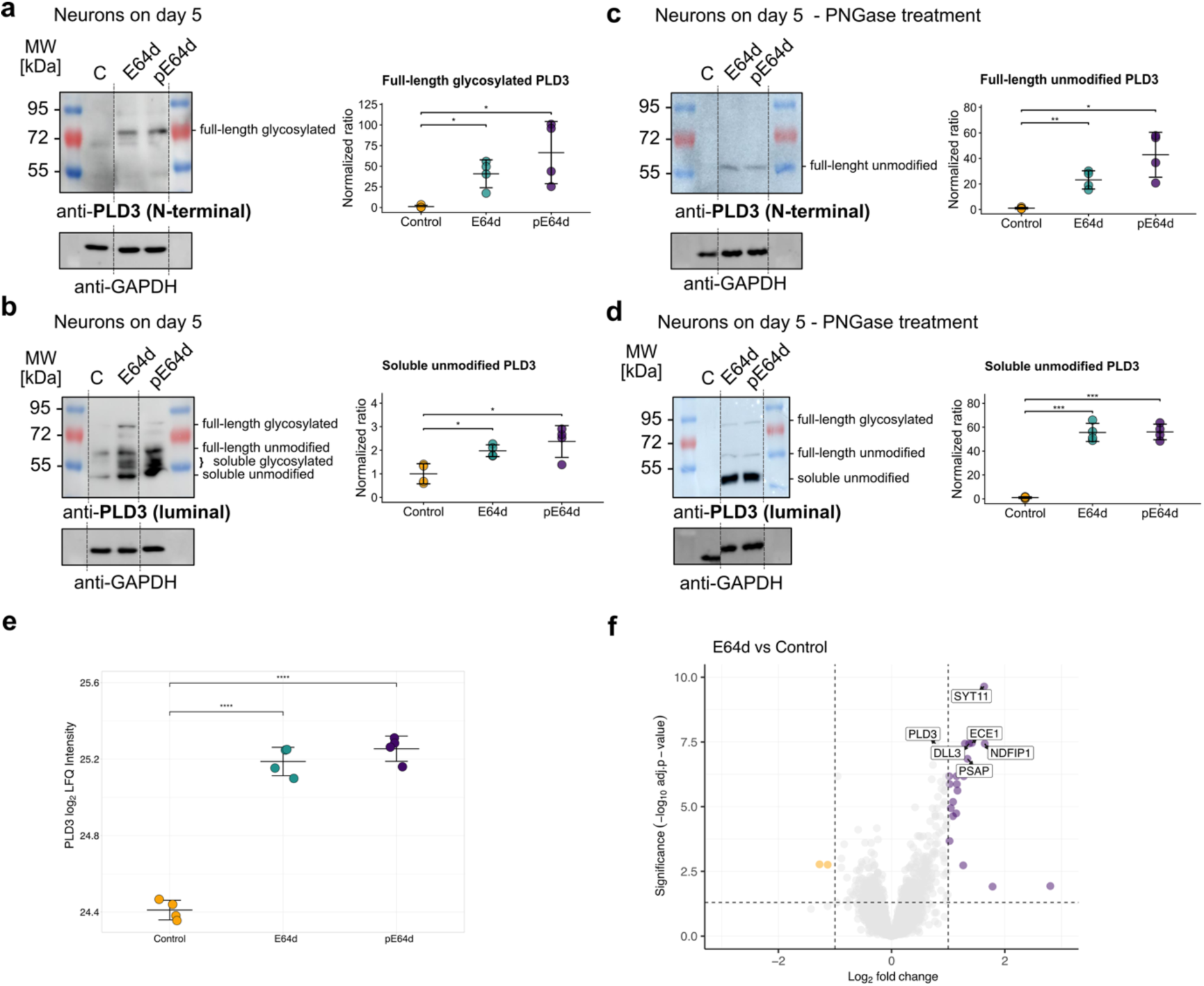
Inhibition of lysosomal cathepsins by E64d leads to accumulation of soluble PLD3 in neurons. **a**) Left, western blot of 5-day iNGN neurons treated with E64d or pE64d using anti-PLD3 (N-terminal) antibody. Right, quantification of full-length glycosylated PLD3. **b**) Left, western blot of 5-day iNGN neurons treated with E64d or pE64d using anti-PLD3 (luminal) antibody. Right, quantification of soluble unmodified PLD3. **c**-**d**) Left, western blots of PLD3 using the N-terminal (**c**) or luminal (**d**) antibody. Right, corresponding quantifications of full-length unmodified (**c**) and soluble unmodified (**d**) PLD3. See also Figure S17. **e**) Dot plot showing the quantification of PLD3 in 5-day iNGN neurons by LC-MS/MS proteomics. (*n* = 4 biological replicates) **f**) Volcano plot visualizing the changes of the whole proteome determined by LC-MS/MS proteomics. Orange dots depict the downregulated proteins. Purple dots show the upregulated proteins; (*n* = 4 biological replicates, cut-off p-value < 0.05, and fold-change > 1). **a**-**d**) The line is mean ± s.d. of *n* = 4 biological replicates. * *p* < 0.05, ** *p* < 0.01, *** *p* < 10^-3^ and **** *p* < 10^-4^, determined by two sample *t*-test. See Figure S21 for replicates. For quantification, the protein intensity was normalized to the loading control and then to the experimental control. In the case of no visible band in the loading control, the mean value of other bands of the same group was used.

Overall, for both E64d and pE64d treatment there was only a minor change in the proteome (**Figure 6f**, see Figure S18 for pE64d). From a total of 7,564 identified proteins, only 21 proteins were significantly upregulated after the treatment with E64d and 15 proteins after addition of pE64d with overlap of eleven proteins (Figure S18-20). Although there was no strong dysregulation of protein degradation on the proteome level, proteins which showed increased levels upon E64d treatment included synaptotagmin-11 (SYT11), Endothelin-converting enzyme 1 (ECE1), NEDD4 family interacting protein 1 (NDFIP1), delta-like protein 3 (DLL3) and mitochondrial carrier homolog 1 (PSAP), pointing towards processes associated with vesicle trafficking^[43]^, neuronal differentiation^[44]^, and apoptosis^[45]^. The analysis of the whole proteome changes showed only negligible number of downregulated proteins. Together, combination of activity-based profiling of cysteine cathepsins in neurons, western blot analysis of PLD3 proteoforms and LC-MS/MS proteomics demonstrate that sole inhibition of cysteine cathepsins (CTSB, CTSC and CTSZ) by E64d leads only to minor proteome changes, but significant accumulation of full-length and N-glycosylated PLD3 proteoforms.

## Conclusions

In summary, we developed a chemical proteomics probe based on E64d that enabled comprehensive profiling of its cellular target landscape and cysteine reactivity. Our results confirm robust engagement of cysteine cathepsins, particularly CTSB, while also revealing previously unrecognized off-target proteins, including BLMH, SUPT5H, PSMA1, KEAP1, and IRAK1, thereby expanding the known interactome of this widely used inhibitor. Despite identifying these additional targets, we found no evidence for alternative lysosomal proteases that could account for PLD3 processing or degradation. Interestingly, the inhibition of cysteine cathepsins in neurons lead to significant accumulation of PLD3 and its dysregulated processing. Given that in patients with Alzheimer’s disease the PLD3 levels are decreased^[17,20,26]^, our findings suggest that this might be caused by activation of cysteine cathepsins in neurons. However, this is contradictory to observations linking neurodegenerative diseases in general with lysosomal insufficiency^[2,7]^. Therefore, our results provide a new mechanistic insight into the regulation of PLD3 and underscore the importance of coordinated cysteine cathepsin activity in neuronal lysosomes that needs to be further investigated. Collectively, our work refines the understanding of E64d selectivity, provides a framework for interpreting its biological effects, and establishes the pE64d probe as a versatile tool for investigating lysosomal protease activity and proteostasis in the context of neurodegeneration.

## Acknowledgements

We are grateful for the support from the Deutsche Forschungsgemeinschaft (DFG, German Research Foundation) with funds from SFB1309 (Chemical Biology of Epigenetic Modifications), project 325871075 (P. K.); M.H. and P.K. were further supported by the BMFTR in the framework of the Cluster4Future program (Cluster for Nucleic Acid Therapeutics Munich, CNATM) (Project ID: 03ZU1201AA) and Boehringer Ingelheim Foundation – Plus 3 Program.

## Data availability

Supporting Information (PDF) contains supporting figures, detail description of the organic synthesis, biochemical methods, proteomic methods and list of reagents used in the study.

All MS data are available via ProteomeXchange with identifier PXD081252.

**Token:** 3zp7tmwzoMZs

