## Supporting Information for "Activity-based chemical proteomics uncovers unexpected covalent targets of E64d and reveals a role for cysteine cathepsins in PLD3 proteostasis"

### Supplementary Figures

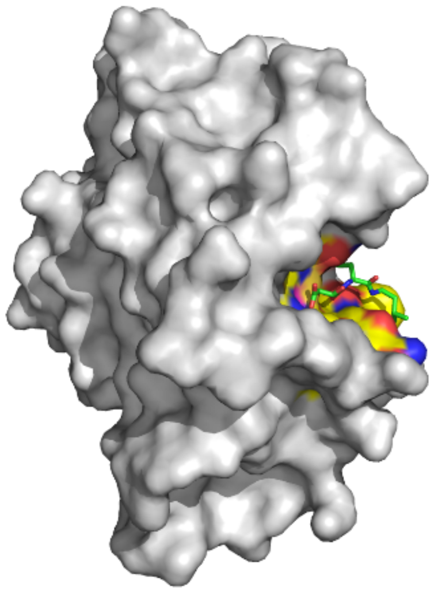

**Figure S1. Crystal structure of bovine Cathepsin B with E64c, the synthetic free acid analogue of E64d.** PDB 1ITO[1]. E64c engages a pocket on the surface of CTSB and the C2 position is covalently bound by the catalytic cysteine residue of CTSB. The aliphatic side chain of the amide bond faces towards the solvent, allowing for modification on this position without loss of CTSB engagement. The leucine residue forms lipophilic interactions with the protein surface (red area above the residue).

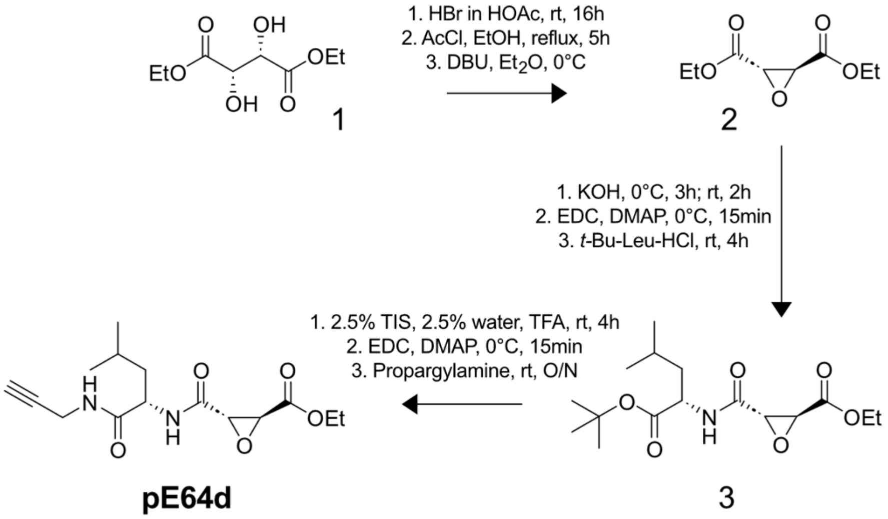

**Figure S2. Synthesis of pE64d.**

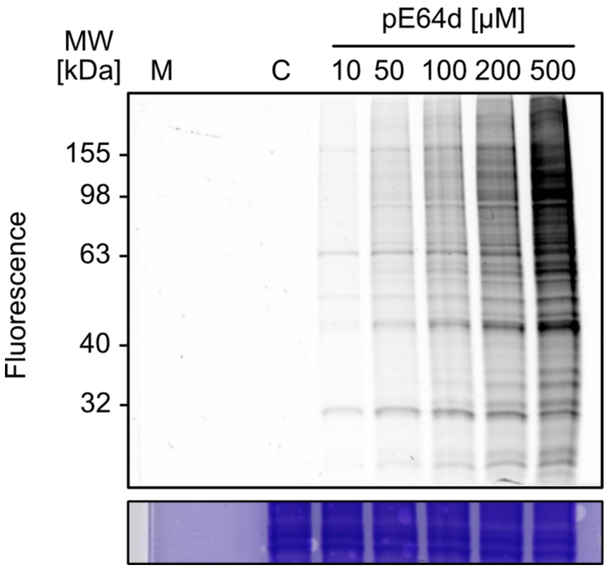

**Figure S3. Concentration-dependent protein labelling by pE64d in HEK293T cells.** HEK293T cells were treated with the indicated concentrations of pE64d or DMSO as a control, incubated for 4 h, lysed and then subjected to CuAAC with TAMRA-$N_3$. For each sample 20 µg protein was loaded onto a 10 % SDS gel.

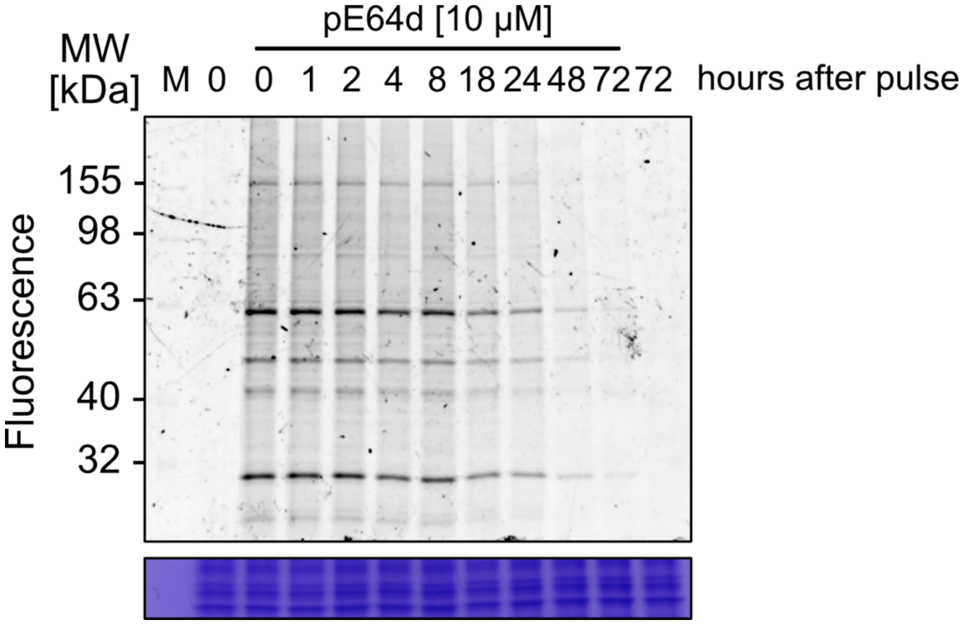

**Figure S4. Time-resolved de-labelling of pE64d-labelled proteins**. HEK293T cells were treated with 10 µM pE64d or DMSO as a control and incubated for 4 h. Culture medium was exchanged to probe-free medium, and cells were incubated for the indicated time before they were harvested. Zero hours after pulse means cells were harvested before the medium was changed.

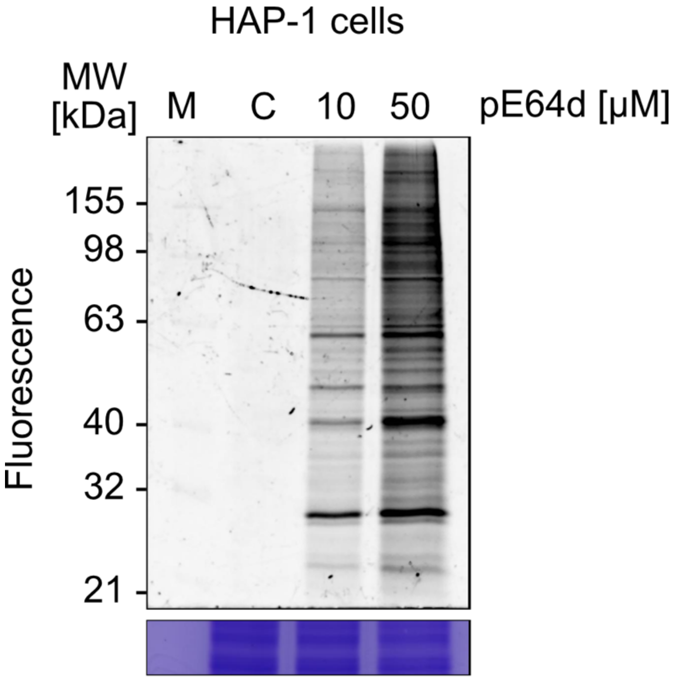

**Figure S5. Concentration-dependent labelling in HAP-1 cells.** HAP-1 cells were treated with 10 µM or 50 µM pE64d or DMSO as a control, incubated for 4 h, lysed and then subjected to CuAAC with TAMRA-$N_3$. For each sample 20 µg protein was loaded onto a 10 % SDS gel.

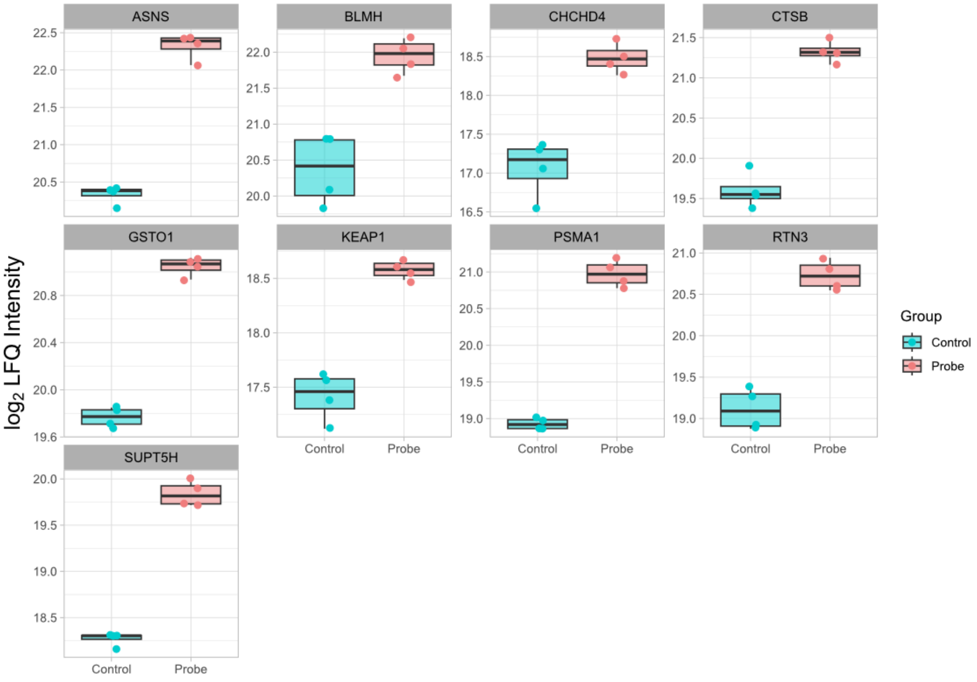

**Figure S6. Profile plots of significantly enriched proteins using 10 µM pE64d.** Protein levels are represented by their $log_2$ LFQ intensity (n = 4 biological replicates).

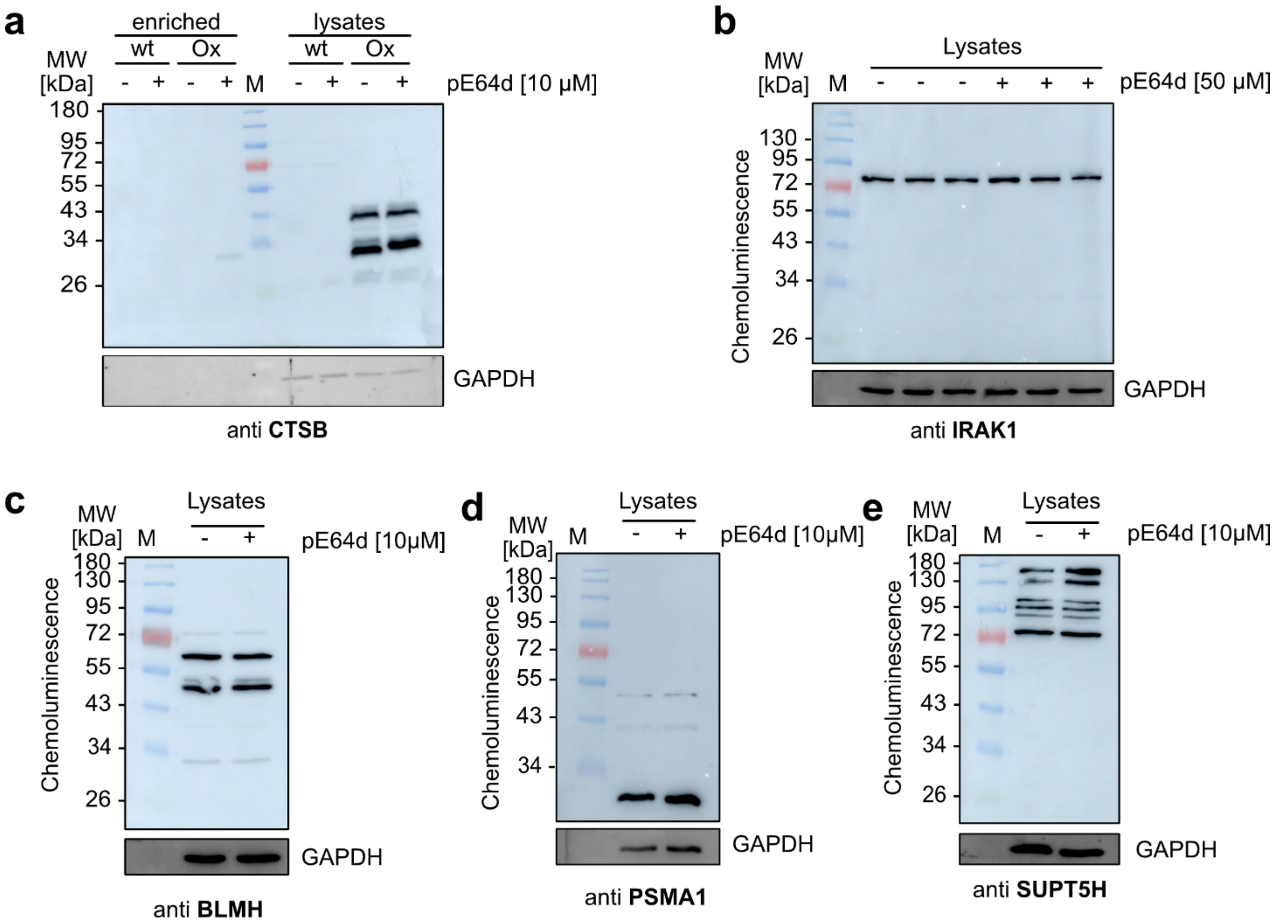

**Figure S7. Whole cell lysate western blot analysis of pE64d target-proteins.** HEK239T cell lysates were subjected to SDS PAGE and subsequent western blot without prior enrichment to verify equal protein levels in both the pE64d treated and DMSO (control) treated lysates. For each sample 20 µg protein was used. Western blot analysis was performed using the following antibodies: **(a)** anti Cathepsin B (with both enriched and non-enriched samples loaded on the same gel), **(b)** anti IRAK1 (biological triplicates), **(c)** anti BLMH, **(d)** anti PSMA1, and **(e)** anti SUPT5H

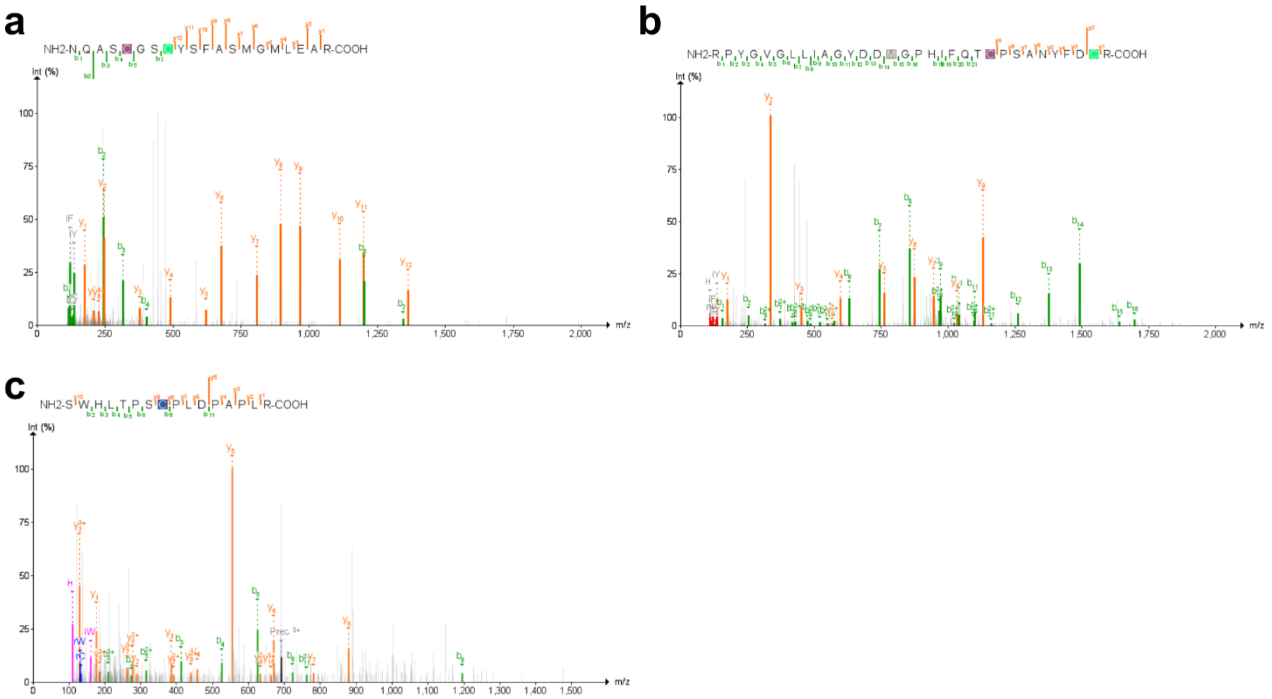

**Figure S8. Assigned MS$^2$ spectra from site-ID experiments. (a)** Cathepsin C, **(b)** PSMA1 and **(c)** IRAK1. Raw files were searched with FragPipe and spectra were visualized by PDV viewer.

D1

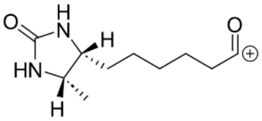

Chemical Formula: $C_{10}H_{17}N_2O_2^+$
Exact Mass: 197,12845

D2

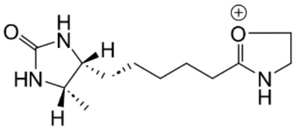

Chemical Formula: $C_{12}H_{22}N_3O_2^+$
Exact Mass: 240,17065

D3

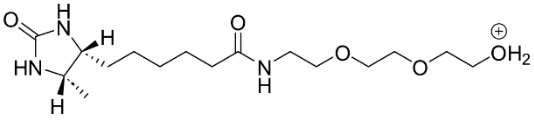

Chemical Formula: $C_{16}H_{32}N_3O_5^+$
Exact Mass: 346,23365

D4.1

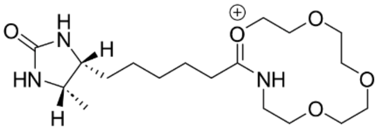

Chemical Formula: $C_{18}H_{34}N_3O_5^+$
Exact Mass: 372,24930

or

D4.2

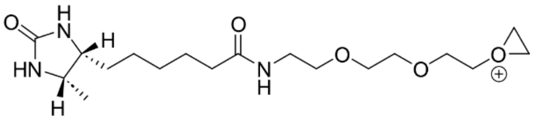

Chemical Formula: $C_{18}H_{34}N_3O_5^+$
Exact Mass: 372,24930

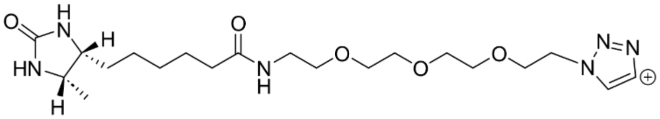

Chemical Formula: $C_{20}H_{35}N_6O_5^+$
Exact Mass: 439,26634

- $N_2$
+ 4 H

D6.1

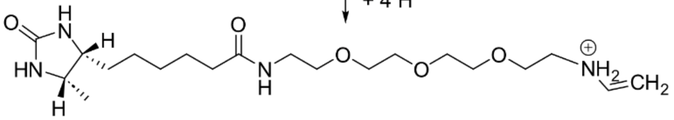

Chemical Formula: $C_{20}H_{39}N_4O_5^+$
Exact Mass: 415,29150

D6.2

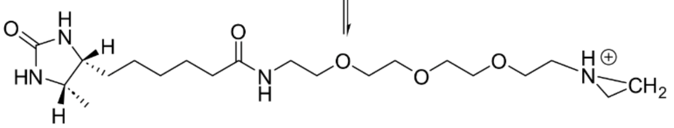

Chemical Formula: $C_{20}H_{39}N_4O_5^+$
Exact Mass: 415,29150

- $C_2H_2$

D5

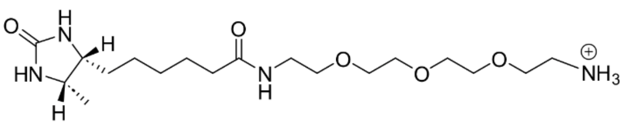

Chemical Formula: $C_{18}H_{37}N_4O_5^+$
Exact Mass: 389,27585

D9

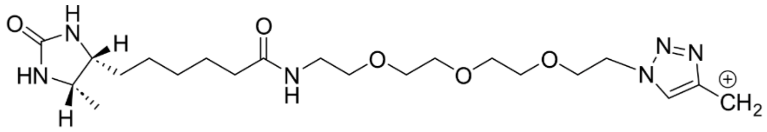

Chemical Formula: $C_{21}H_{37}N_6O_5^+$
Exact Mass: 453,28199

↓ - $N_2$

D7

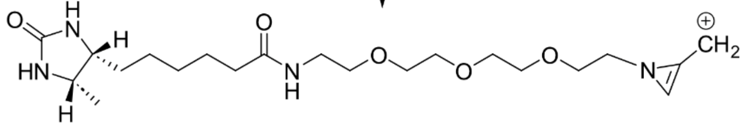

Chemical Formula: $C_{21}H_{37}N_4O_5^+$
Exact Mass: 425,27585

D10

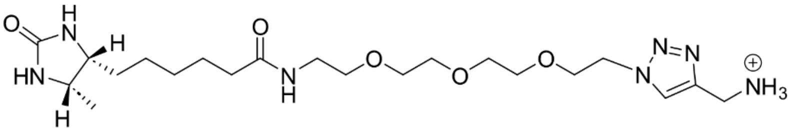

Chemical Formula: $C_{21}H_{40}N_7O_5^+$
Exact Mass: 470,30854

↓ - $N_2$

D8

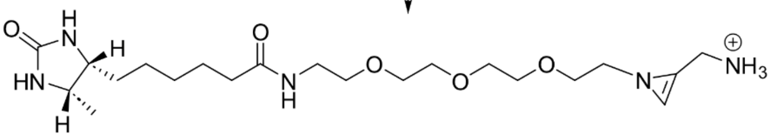

Chemical Formula: $C_{21}H_{40}N_5O_5^+$
Exact Mass: 442,30240

**D11**

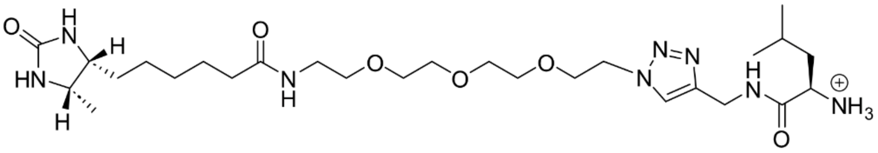

Chemical Formula: $C_{27}H_{51}N_8O_6^+$
Exact Mass: 583,39261

**D12**

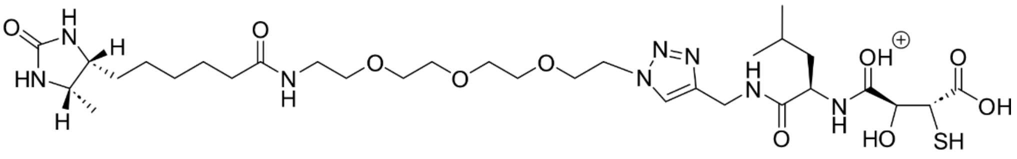

Chemical Formula: $C_{31}H_{55}N_8O_{10}S^+$
Exact Mass: 731,37564

**Figure S9. Diagnostic ions from Desthiobiotin-E64d-modified peptides**. Raw LC-MS/MS data were searched in FragPipe using a custom diagnostic ion mining workflow. Diagnostic ion masses were extracted from the “mass” column of the global.diagmine output file. These masses were matched to possible structures. The identified diagnostic ions were designated as D1 through D12 in ascending order of their mass-to-charge ratios (m/z). D11 and D12 represent pE64d specific diagnostic ions.

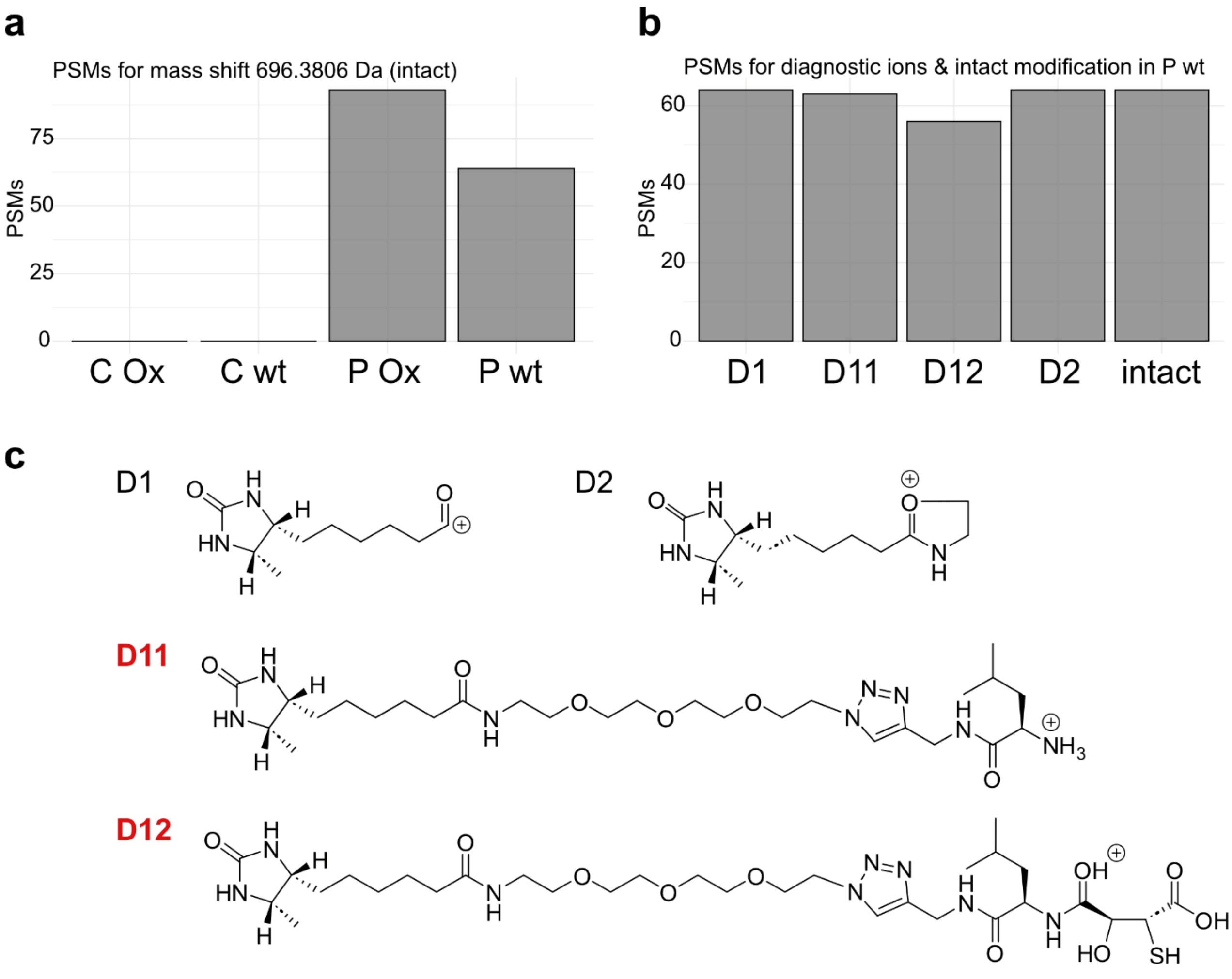

**Figure S10. Diagnostic ion PSMs. (a)** Number of PSMs for the intact modification (+ 696.3806 Da) across all samples. Overall, 93 PSMs were observed in the pE64d-treated group with CTSB wild-type overexpression (P Ox), whereas 64 PSMs were observed in the pE64d-treated group with no overexpression of CTSB (P wt). No PSMs were observed in either control group. **(b)** Number of PSMs for diagnostic ions D1, D2, D11, D12, and the intact modification in sample P wt. Diagnostic ions D1 and D2 were observed in all 64 PSMs containing the intact modification. Diagnostic ion D11 was present in 63, and diagnostic ion D12 was present in 56 of these PSMs, respectively. **(c)** Structures of the discussed diagnostic ions.

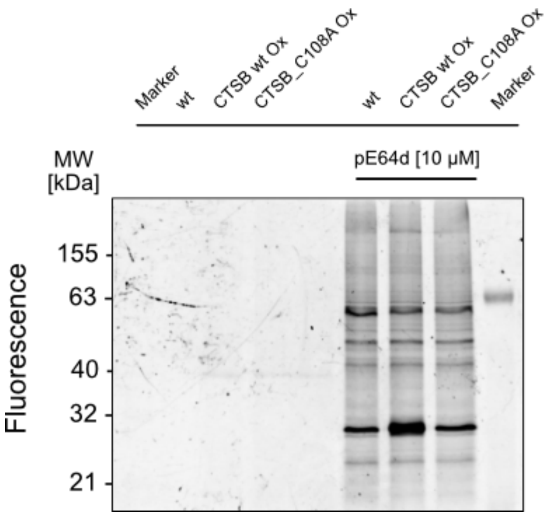

**Figure S11. In-gel fluorescence analysis of pE64d treated wild type and transfected HEK293T cells.** Lysates from pE64d (10 µM) treated wild type HEK293T cells or HEK293T cells overexpressing either wild-type CTSB or the CTSB C108A mutant were subjected to CuAAC with TAMRA-Biotin-$N_3$ and subsequently enriched using the SP2E protocol. Figure shows the in-gel fluorescence of the SDS PAGE.

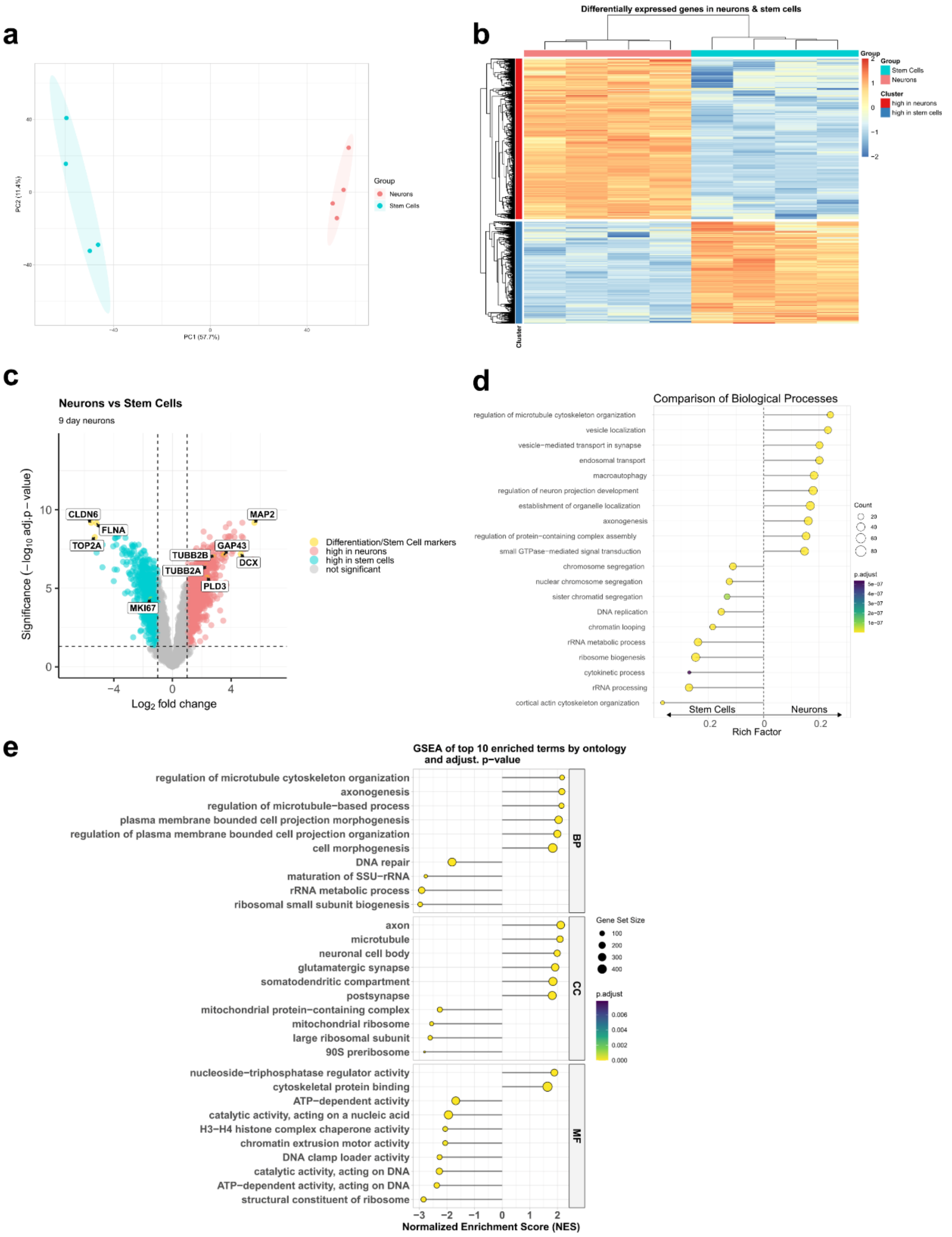
a
Group
Neurons
Stem Cells
PC2 (11.4%)
PC1 (57.7%)
b
Differentially expressed genes in neurons & stem cells
Group
Stem Cells
Neurons
Cluster
high in neurons
high in stem cells
c
Neurons vs Stem Cells
9 day neurons
Significance (−log10 adj.p − value)
Log2 fold change
CLDN6
FLNA
TOP2A
MAP2
GAP43
TUBB2B
DCX
TUBB2A
PLD3
MKI67
Differentiation/Stem Cell markers
high in neurons
high in stem cells
not significant
d
Comparison of Biological Processes
regulation of microtubule cytoskeleton organization
vesicle localization
vesicle−mediated transport in synapse
endosomal transport
macroautophagy
regulation of neuron projection development
establishment of organelle localization
axonogenesis
regulation of protein−containing complex assembly
small GTPase−mediated signal transduction
chromosome segregation
nuclear chromosome segregation
sister chromatid segregation
DNA replication
chromatin looping
rRNA metabolic process
ribosome biogenesis
cytokinetic process
rRNA processing
cortical actin cytoskeleton organization
Count
p.adjust
Stem Cells
Neurons
Rich Factor
e
GSEA of top 10 enriched terms by ontology and adjust. p−value
regulation of microtubule cytoskeleton organization
axonogenesis
regulation of microtubule−based process
plasma membrane bounded cell projection morphogenesis
regulation of plasma membrane bounded cell projection organization
cell morphogenesis
DNA repair
maturation of SSU−rRNA
rRNA metabolic process
ribosomal small subunit biogenesis
axon
microtubule
neuronal cell body
glutamatergic synapse
somatodendritic compartment
postsynapse
mitochondrial protein−containing complex
mitochondrial ribosome
large ribosomal subunit
90S preribosome
nucleoside−triphosphatase regulator activity
cytoskeletal protein binding
ATP−dependent activity
catalytic activity, acting on a nucleic acid
H3−H4 histone complex chaperone activity
chromatin extrusion motor activity
DNA clamp loader activity
catalytic activity, acting on DNA
ATP−dependent activity, acting on DNA
structural constituent of ribosome
BP
CC
MF
Gene Set Size
p.adjust
Normalized Enrichment Score (NES)

**Figure S12. Verification of successful differentiation of iNGNs into neurons**. Whole proteome was measured from stem cells and neurons (n = 4 biological replicates for each group). **(a)** PCA plot of stem cells and neurons. **(b)** Heatmap showing differentially expressed genes. **(c)** Volcano plot of differentially expressed proteins from stem cells and neurons. Differentiation or proliferation markers are highlighted in yellow. $Log_2$ fold change cutoff > 1, $-log_{10}$ (adj.p-value) cutoff < 0.05. **(d)** GO term enrichment analysis of biological processes of stem cells and neurons. **(e)** Gene set enrichment analysis (GSEA) representing the ten most significant terms by adjusted p-value. Ontologies are biological process (BP), cellular component (CC) and molecular function (MF). Terms with positive NES are enriched in neurons, terms with negative NES are enriched in stem cells.

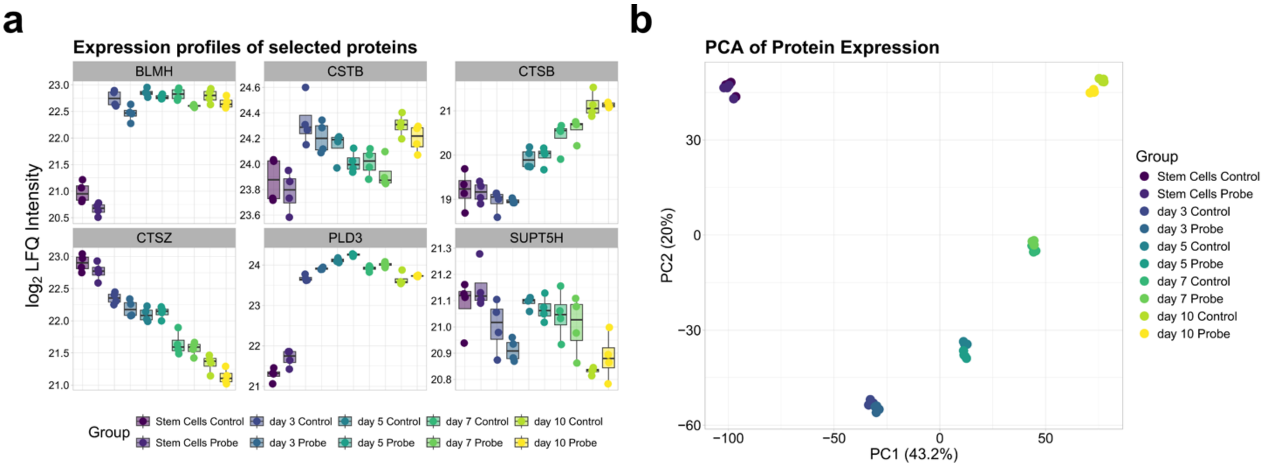

**Figure S13. Whole proteome analysis of hiPSCs and neurons during differentiation. (a)** Protein levels of stem cells and neurons on different time-points of neuronal differentiation. **(b)** PCA of protein expression of stem cells and neurons on different time-points of neuronal differentiation.

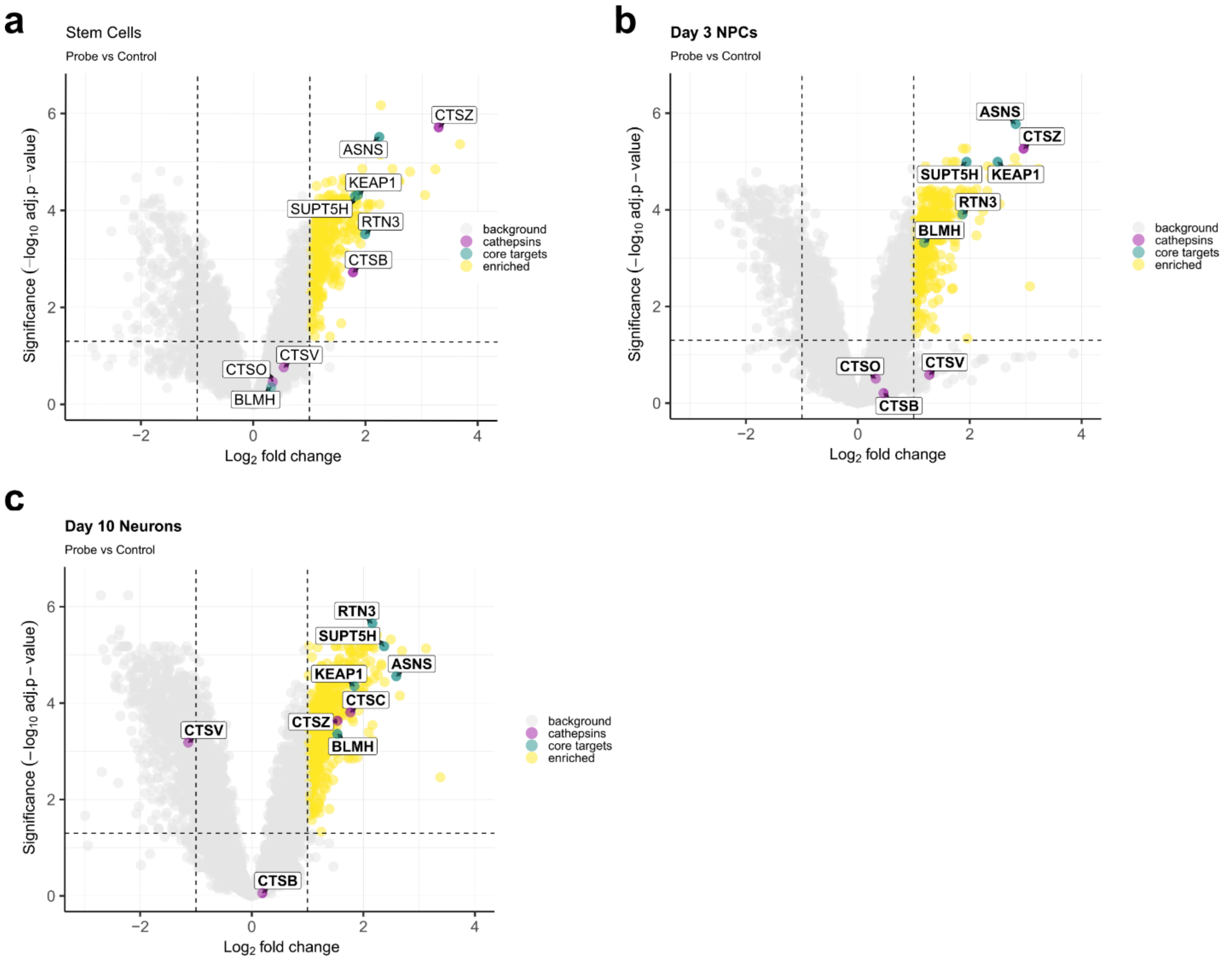

**Figure S14. Volcano plots of pE64d-enriched proteins.** Differential enrichment is shown for **(a)** stem cells, **(b)** 3-day-old neurons and **(c)** 10-day-old neurons. Significantly enriched proteins are represented by yellow circles (n = 4 technical replicates, cut-off p-value < 0.05, and fold-change > 1). Cysteine cathepsins are depicted in magenta. The group of the most significantly enriched proteins across various conditions is depicted in green.

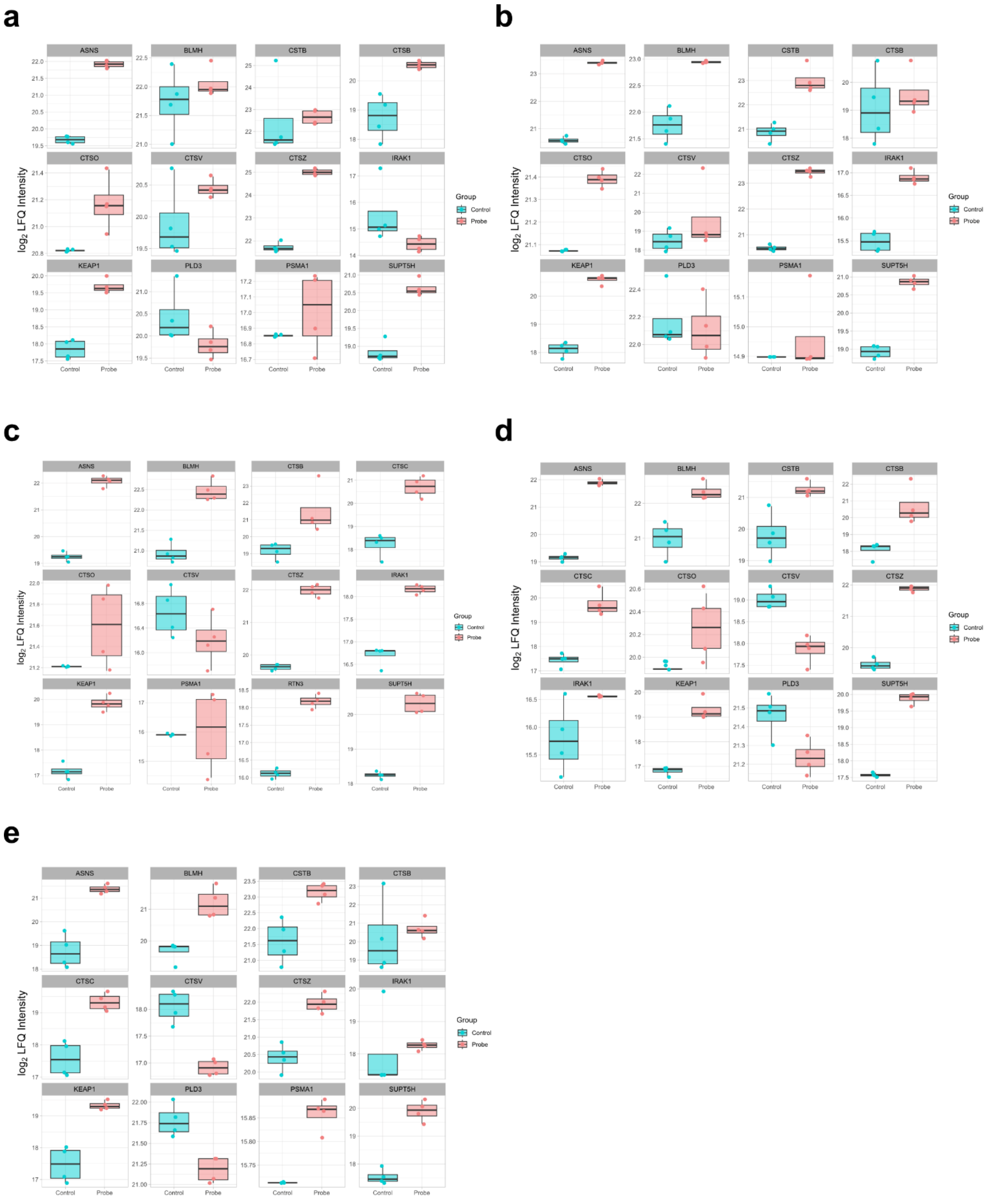

**Figure S15. Profile plots of selected proteins from pE64d- (50 µM) treated neurons or stem cells after enrichment by SP2E.** Protein levels are represented by their $\log_2$ LFQ intensity. Profiles are shown for **(a)** stem cells; **(b)** neurons on day 3; **(c)** neurons on day 5; **(d)** neurons on day 7, and **(e)** neurons on day 10 of differentiation.

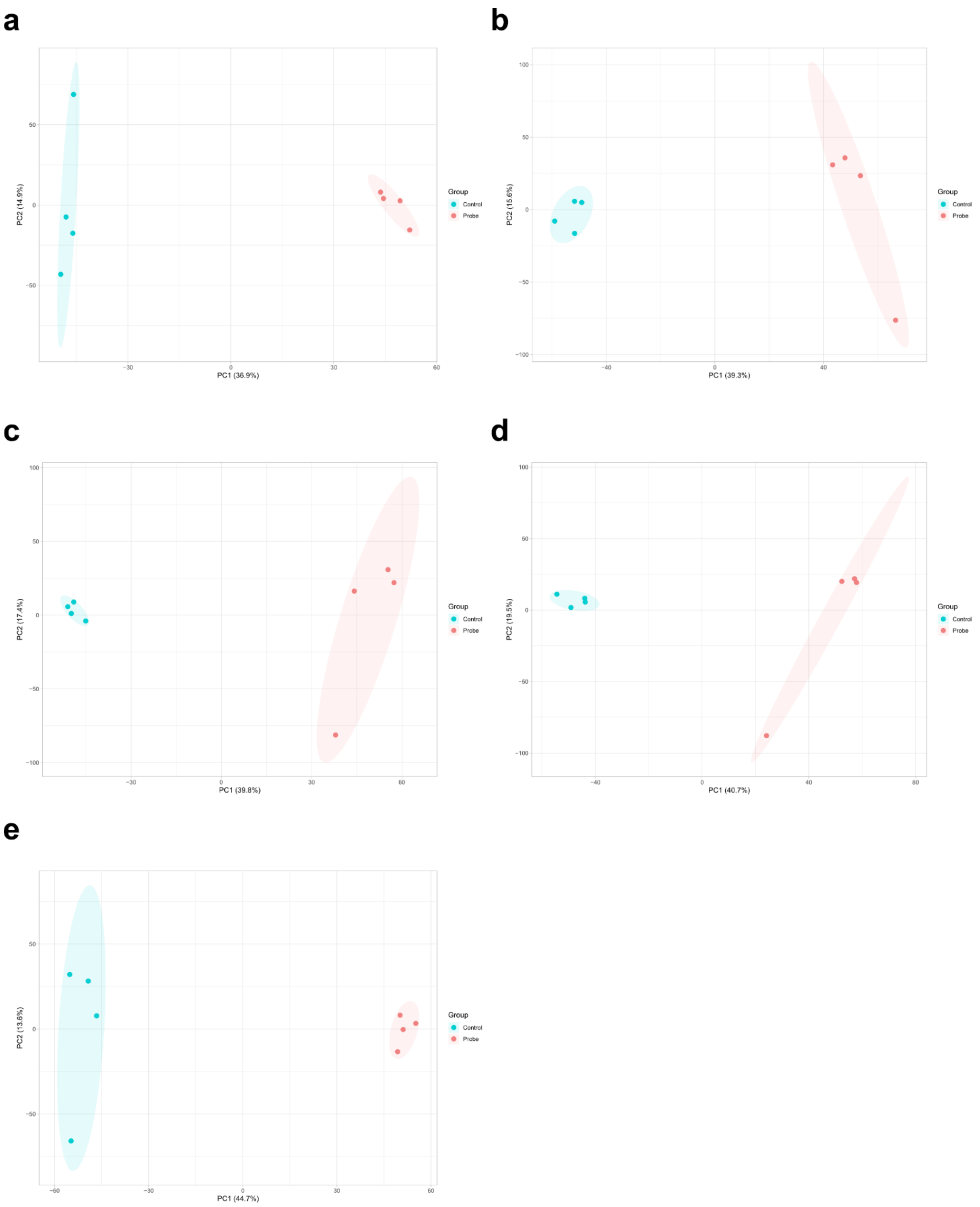

**Figure S16. PCA plots of pE64d- (50 µM) or DMSO-treated neurons or stem cells after enrichment. (a)** Stem cells; **(b)** neurons on day 3; **(c)** neurons on day 5; **(d)** neurons on day 7; and **(e)** neurons on day 10 of differentiation.

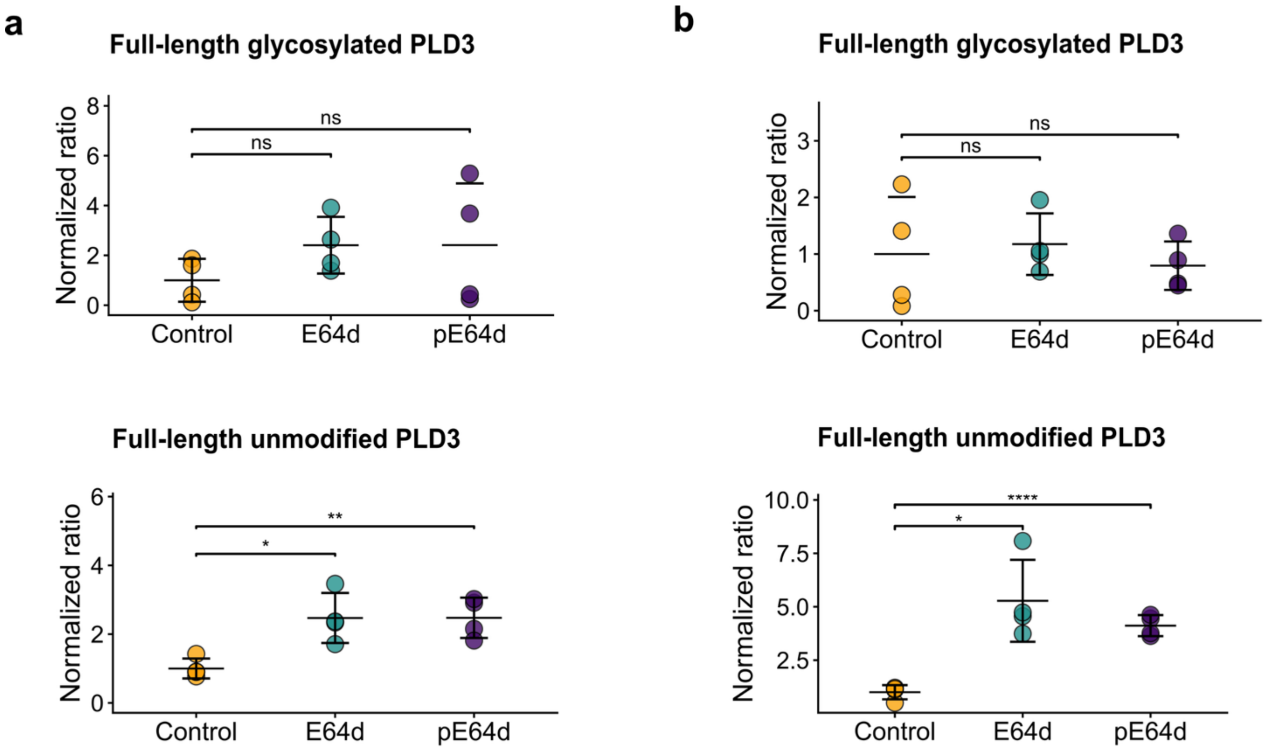

**Figure S17**. **Quantification of PLD3 forms of 5-day iNGN neurons treated with E64d or pE64d using anti-PLD3 (luminal) antibody (a)** before and **(b)** after PNGase F treatment.

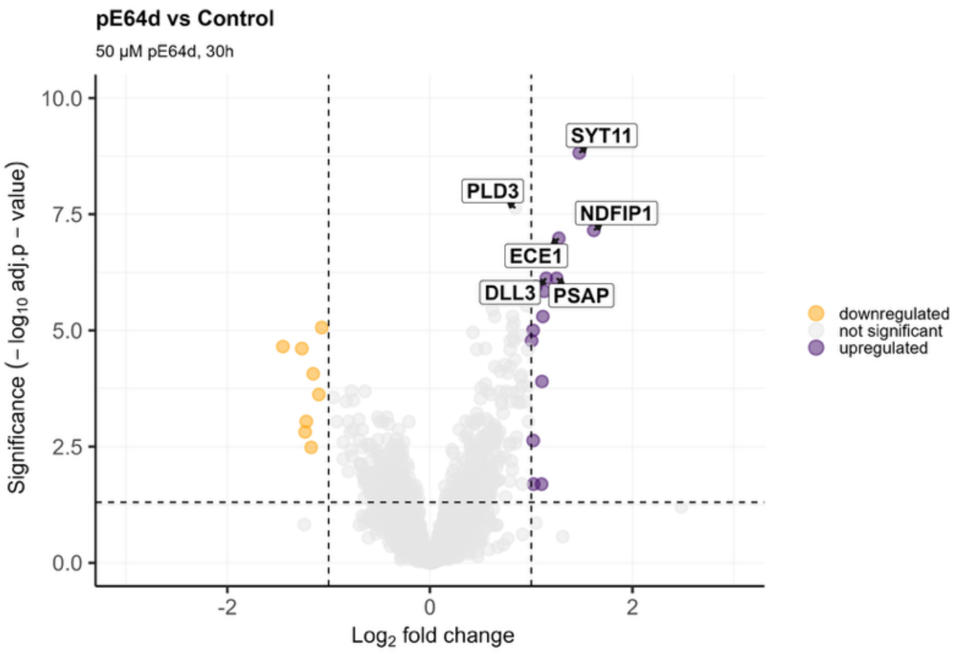

**Figure S18. Differentially expressed proteins after pE64d treatment.** Volcano plot visualizing the changes of the whole proteome determined by LC-MS/MS proteomics. Orange dots depict the downregulated proteins. Magenta dots show the upregulated proteins; ($n$ = 4 biological replicates, cut-off $p$-value < 0.05, and fold-change > 1).

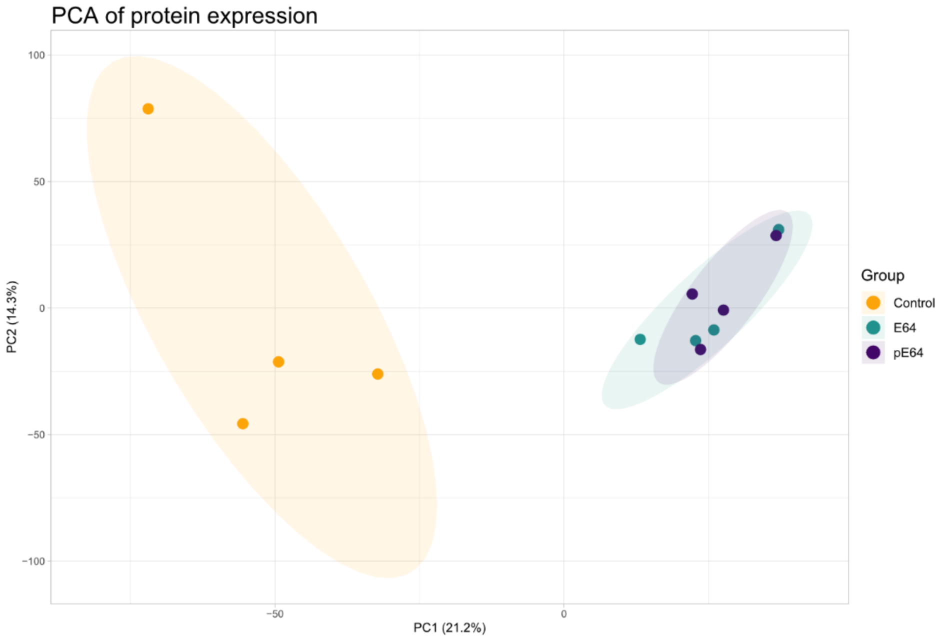

**Figure S19. Principal component analysis of whole proteome analysis of control, E64d and pE64d treated day 5 iNGN neurons.**

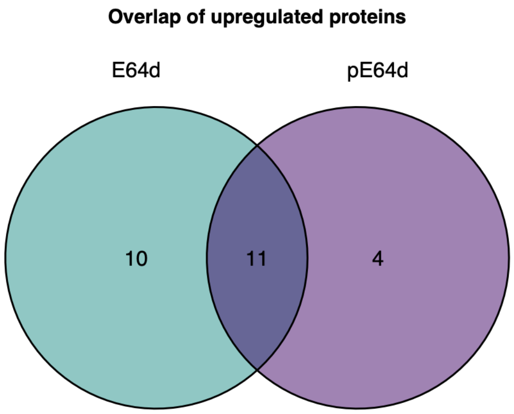

**Figure S20. Overlap between significantly upregulated proteins upon E64d and pE64d treatment in 5-day iNGN neurons.**

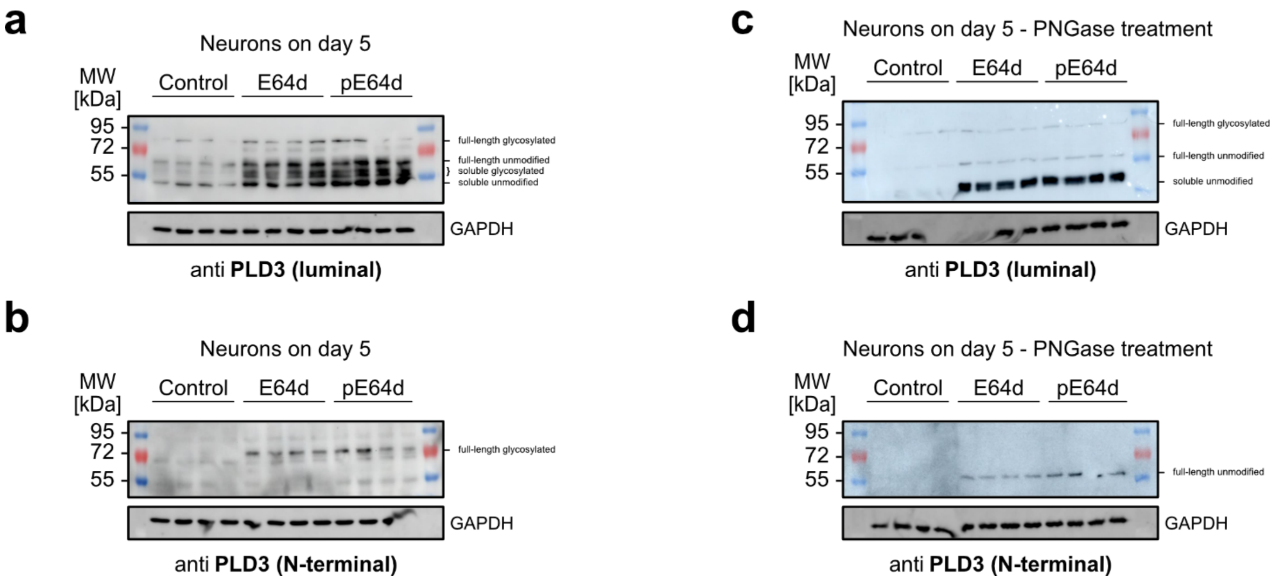

**Figure S21. Western blots representing all four biological replicates of main figure 6. a**) Western blot of 5-day iNGN neurons treated with E64d or pE64d using anti-PLD3 (luminal) antibody. **b**) Western blot of 5-day iNGN neurons treated with E64d or pE64d using anti-PLD3 (N-terminal) antibody. **c**-**d**) Western blots of PLD3 using the luminal (**c**) or N-terminal (**d**) antibody after PNGase treatment. For quantification, the protein intensity was normalized to the loading control and then to the experimental control. In the case of no visible band in the loading control, the mean value of other bands of the same group was used.

**Supplementary Tables**

**Table S1**. PSMs of indicated mass shifts (Peak column) and diagnostic ions in sample P wt from the Site-ID experiment.

| Peak | PSMs | 197.12845 PSMs | 240.17065 PSMs | 583.39261 PSMs | 731.37564 PSMs |
|---|---|---|---|---|---|
| **0.0006** | 81 | 64 | 56 | 21 | 3 |
| **696.3806** | 64 | 64 | 64 | 63 | 56 |
| **57.0206** | 4 | 3 | 3 | 2 | 0 |
| **0.0112** | 7 | 4 | 4 | 1 | 0 |
| **-0.0114** | 5 | 5 | 5 | 3 | 3 |

**Table S2.** Detected diagnostic ions in the diagnostic ion mining search (FragPipe)

| Diagnostic Ion | Chemical Formula (for charge +1) | Calculated Mass [Da] | Found Mass [Da] | Delta Mass | Percent in modified PSMs |
|---|---|---|---|---|---|
| **D1** | C10H17N2O2 | 197.12845 | 197.1282 | -0.00025 | 100 |
| **D2** | C12H22N3O2 | 240.17065 | 240.1704 | -0.00025 | 100 |
| **D3** | C16H32N3O5 | 346.23365 | 346.2336 | -5E-05 | 59.26 |
| **D4** | C18H34N3O5 | 372.2493 | 372.2488 | -0.0005 | 25.93 |
| **D5** | C18H37N4O5 | 389.27585 | 389.2756 | -0.00025 | 100 |
| **D6** | C20H39N4O5 | 415.2915 | 415.2914 | -1E-04 | 72.22 |
| **D7** | C21H37N4O5 | 425.27585 | 425.2756 | -0.00025 | 100 |
| **D8** | C21H40N5O5 | 442.3024 | 442.3022 | -0.0002 | 100 |
| **D9** | C21H37N6O5 | 453.282 | 453.282 | 0 | 100 |
| **D10** | C21H40N7O5 | 470.30854 | 470.3084 | -0.00014 | 100 |
| **D11** | C27H51N8O6 | 583.39261 | 583.3926 | -1E-05 | 100 |
| **D12** | C31H55N8O10S | 731.37564 | 731.3762 | 0.00056 | 77.78 |
| **D2** | C12H23N3O2 (charge +2) | 120.5889 | 120.589 | 0.0001 | 25.93 |

**Table S3.** Conditions for PCR cycle. The annealing temperature $T_a$ for each primer pair was determined based on its sequence.

| Step | Temperature | Time |
|---|---|---|
| Initial Denaturation | 98 °C | 30 seconds |
| 25 Cycles | 98 °C | 10 seconds |
| | $T_a$ | 30 seconds |
| | 72 °C | 30 seconds/kb |
| Final Extension | 72 °C | 2 minutes |
| Hold | 4 °C | |

**Table S4.** Sequences of oligonucleotides (5' to 3') used for sequencing.

| Construct | Sense | Sequence (5' to 3') |
|---|---|---|
| CTSB C108A | Forward | ACACCAATGCGCACGTCA |

**Table S5.** Sequences of oligonucleotides (5' to 3') used for cloning. $T_a$ - annealing temperature.

| Protein | Mutant | Sense | Sequence (5' to 3') | $T_a$ [°C] |
|---|---|---|---|---|
| CTSB | C108A | Forward | CTGTGGCAGCGCGTGGGCCTTCG | 83.1 |
| | | Reverse | GAGCCCTGGTCTCTG | 54.4 |

**Organic Synthesis**

Reagents and solvents were purchased from commercial suppliers, including abcr GmbH (Karlsruhe, Germany), Acros Organics (Thermo Fisher Scientific, USA), Sigma-Aldrich (St. Louis, USA), TCI Deutschland GmbH (Eschborn, Germany), Thermo Scientific (Loughborough, UK) and they were used without any further purification.

TLC (thin layer chromatography) was performed to monitor the reaction progress, which was done on precoated silica gel plates (60 F-254, 0.25 mm, from Merck KGaA, Darmstadt, Germany) as the stationary phase, with detection by potassium permanganate staining and gentle heating.

Flash chromatography was performed on Pure Chromatography Instruments (Pure-C815 Flash) from BUCHI Labortechnik GmbH (Essen, Germany) on pre-fabricated FlashPure EcoFlex Silica columns (25 or 40 g) with the indicated eluents.

1H and proton-decoupled 13C NMR spectra for compound characterization in deuterated chloroform ($CDCl_3$) were acquired at 298K on a Bruker Avance Neo 500 spectrometer (11.7 T), using a nitrogencooled Prodigy BBO probe. All chemical shifts are reported in delta (δ) units in parts per million (ppm) relative to distinguished solvent signals as an internal reference. Coupling constant *J* are indicated in Hertz (Hz). Splitting patterns for peak assignments are indicated as the following abbreviations: s, singlet; d, doublet; t, triplet; q, quartet; m, multiplet; dd, doublet of doublets; dt, doublet of triplets. Spectra were analyzed using MestReNova (version 14.2.1, Mestrelab Research, Santiago de Compostela, Spain).

LC-MS chromatograms were obtained from Thermo Fisher LC-MS system composed of a DIONEX UltiMate 3000 HPLC system (pump, auto sampler, column compartment, and diode array detector) and an ESI (electrospray ionization)-MS based MSQ Plus single quadrupole mass spectrometer; the latter one was used for ESI-MS measurements with direct injection. Reversed phase column chromatographic methods use a Hypersil Gold C18 selectivity column (100 × 2.1 mm).

Unless otherwise stated, all reactions were performed under inert atmosphere and water free solvents were used.

**Diethyl (2S,3S)-oxirane-2,3-dicarboxylate**

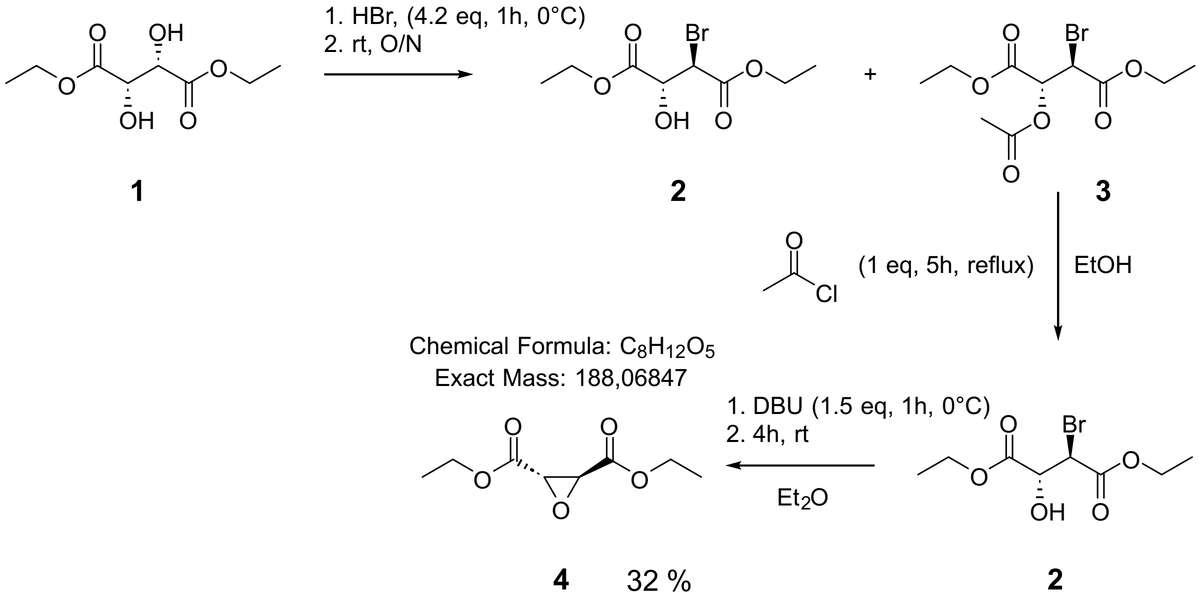

A 30% solution of HBr (4.2 eq, 80 mL, 408 mmol) in HOAc was added at 0 °C to neat (2S, 3S)-Diethyl tartrate **1** (20.0 g, 16.5 mL, 97 mmol) over 1h. Fifteen min after the final HBr addition, the solution was allowed to warm to r.t. and was stirred overnight. The reaction mixture was poured onto crushed ice (200 g), and the resulting mixture was extracted with $Et_2O$ (4× 100 mL). Combined organic layers were washed with $H_2O$ (2x 150 mL) and brine (2x 100 mL), dried over $MgSO_4$ and concentrated under reduced pressure, affording a mixture of the deacetylated and acetylated product **2** and **3** as a pale yellow oil. The crude product was dissolved in anhydrous EtOH (150 mL) and an equimolar amount of acetyl chloride (6.9 mL, 0.097 mol) to **1** was added dropwise. The solution was heated under gentle reflux until TLC analysis indicated complete conversion to the lower running deacetylated product (ca 5 h). After cooling to r.t., the solution was concentrated under reduced pressure to give the bromohydrin **2** as a pale-yellow oil. The crude bromohydrin **2** (17.7 g, 65.8 mmol) was dissolved in anhydrous $Et_2O$ (30 mL) and cooled to 0°C. A solution of DBU (1.5 eq, 14.7 mL, 98.5 mmol) in anhydrous $Et_2O$ (10 mL) was added dropwise for 1 h at 0 °C. After the final addition of DBU, the mixture was allowed to warm to r.t. and stirred for an additional 4 h. The reaction was quenched with cold 1N HCl to pH ~ 5. The resulting aqueous layer was extracted with $Et_2O$ (3x 150 mL). The combined organic fractions were washed with brine (2x 100 mL), dried over $MgSO_4$ and concentrated under reduced pressure to give a faintly yellow oil. Vacuum distillation gave the desired product Diethyl (2S,3S)-oxirane-2,3-dicarboxylate **4** as a colorless oil (5.78 g, 30.7 mmol, 32 %).

$^1$H NMR (500 MHz, $CDCl_3$) δ 4.34 – 4.20 (m, 4H), 3.66 (s, 2H), 1.31 (t, *J* = 7.1 Hz, 6H).

$^{13}$C NMR (126 MHz, $CDCl_3$) δ 166.94, 62.40, 52.18, 14.18.

HR-MS (ESI+): m/z $[M+H]^+$ calculated for $C_8H_{13}O_5$: 189.0757, found: 189.0758

HR-MS (ESI-): m/z $[M-H]^-$ calculated for $C_8H_{11}O_5$: 187.0612, found: 187.0610

**Ethyl (2S,3S)-3-((1-(tert-butoxy)-4-methyl-1-oxopentan-2-yl)carbamoyl)oxirane-2-carboxylate**

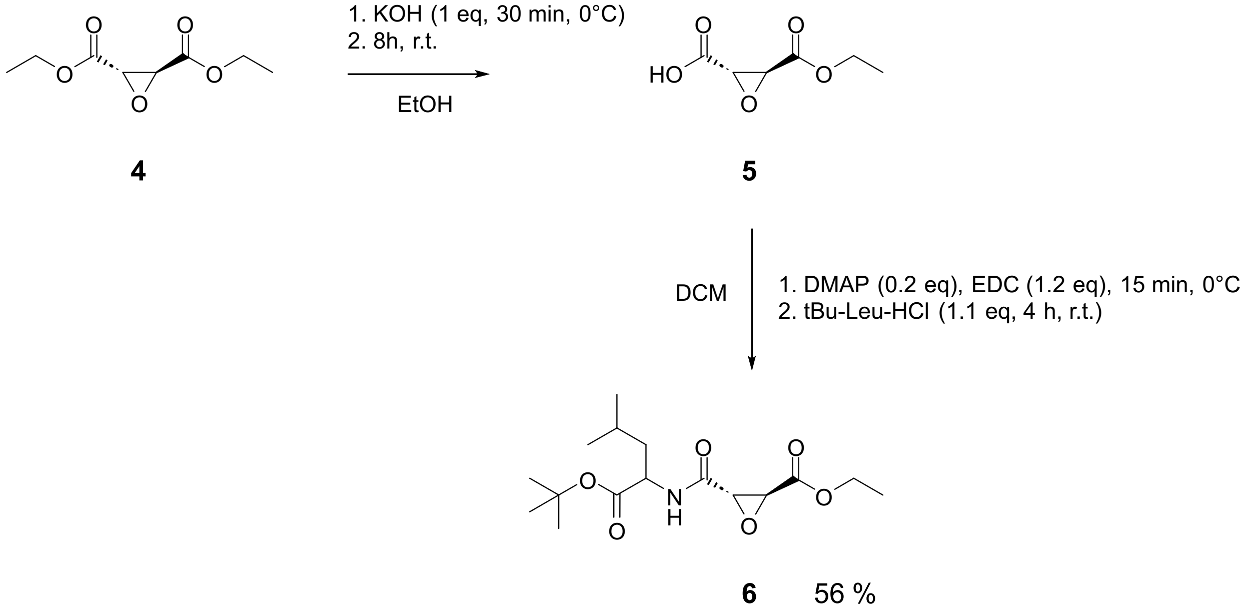

Diethyl (2S,3S)-oxirane-2,3-dicarboxylate **4** (502 mg, 2,67 mmol) was dissolved in absolute EtOH (20 mL) and cooled in an ice-bath. A solution of KOH (1 eq, 150 mg, 2,67 mmol) in absolute EtOH (150 mL) was added dropwise over 15 min. The reaction mixture was stirred at 0 °C for 3 h, and then allowed to warm to r.t. and stirred for an additional 2 h. The solvent was evaporated under reduced pressure, and the resulting white solid was dissolved in $H_2O$ (50 mL). The aqueous solution was washed with DCM (30 mL), acidified to pH 1–2 with conc. HCl, and saturated with NaCl. The mixture was extracted with EtOAc (4 × 100 mL). The combined organic layers were dried over $MgSO_4$ and concentrated to afford the crude monoethyl ester **5** as a colorless oil, which was used in the next step without further purification. To a solution of the crude monoethyl ester **5** in anhydrous DCM (15 mL) at 0 °C were added DMAP (0.2 eq, 65.2 mg, 0.53 mmol) and EDC (1.2 eq, 0.56 mL, 3.2 mmol). The mixture was stirred at 0 °C for 15 min. A suspension of *t*Bu-Leu-HCl (1.1 eq, 657 mg, 2.93 mmol) in DCM (5 mL) was added dropwise, and the reaction was stirred at r.t. for 4 h. The reaction was quenched with $H_2O$, and washed sequentially with 1N HCl, sat. $NaHCO_3$ (aq.) and brine. The organic phase was dried over $MgSO_4$ and concentrated under reduced pressure to afford product **6** (496 mg) which was used in the next step without further purification.

HR-MS (ESI+): m/z $[M+H]^+$ calculated for $C_{16}H_{28}NO_6$: 330.1911, found: 330.1910

**Ethyl (2S,3S)-3-((4-methyl-1-oxo-1-(prop-2-yn-1-ylamino)pentan-2-yl)carbamoyl)oxirane-2-carboxylate**

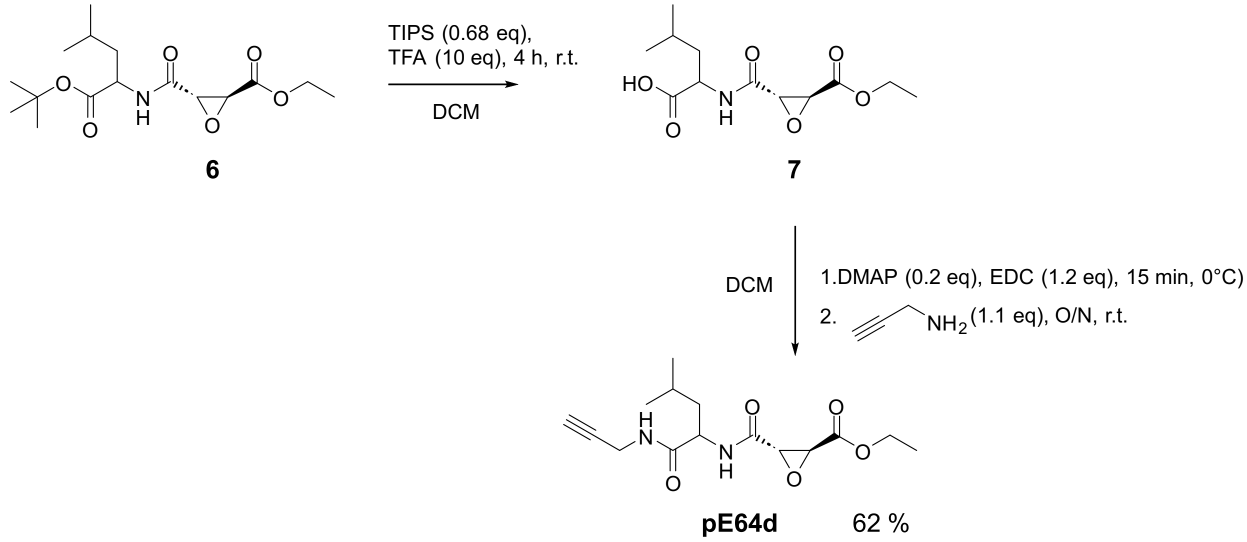

The *tert*-butyl ester **6** (474 mg, 1.44 mmol) was dissolved in DCM (50 mL) and triisopropylsilane (0.68 eq, 0.2 mL) and trifluoroacetic acid (10 eq, 1.1 mL) were added. The colorless solution was stirred for 4 h at r.t., and then concentrated under reduced pressure. Traces of trifluoroacetic acid were removed by repeated co-evaporation with toluene, affording the free carboxylic acid **7** as a viscous pale yellow oil which was used without further purification. To a solution of the carboxylic acid **7** in DCM (15 mL) at 0 °C were added DMAP (0.2 eq, 35.2 mg, 0.29 mmol) and EDC (1.2 eq, 305 µL, 1.73 mmol). The mixture was stirred at 0 °C for 15 min. A solution of propargylamine (1.1 eq, 101 µL, 1.58 mmol) in 5 mL DCM was added dropwise and the reaction was stirred at r.t. overnight. The reaction was quenched with 1N HCl to pH ~ 5 and washed sequentially with 1N HCl, sat. $NaHCO_3$ (aq.), $H_2O$ and brine. The organic phase was dried over $MgSO_4$ and concentrated under reduced pressure. The residue was purified by silica column chromatography (1. Hexane, 200 mL; 2. Hexane : EtOAc 10:1, 300 + 30 mL; 3. Hexane : EtOAc 5:1, 500 + 100 mL) to afford **pE64d** (275 mg, 0.886 mmol, 62 %).

$^{1}H$ NMR (500 MHz, $CDCl_3$) δ 6.53 (d, $J$ = 8.7 Hz, 1H), 6.35 (t, $J$ = 5.4 Hz, 1H), 4.43 (td, $J$ = 8.6, 5.8 Hz, 1H), 4.34 – 4.20 (m, 2H), 4.11 – 3.98 (m, 2H), 3.71 (d, $J$ = 1.8 Hz, 1H), 2.25 (t, $J$ = 2.6 Hz, 1H), 1.73 – 1.63 (m, 1H), 1.61 – 1.49 (m, 2H), 1.32 (t, $J$ = 7.1 Hz, 3H), 0.93 (dd, $J$ = 10.6, 6.3 Hz, 6H).

$^{13}C$ NMR (126 MHz, $CDCl_3$) δ 170.76, 166.52, 166.40, 79.02, 72.17, 62.57, 53.91, 53.16, 51.18, 41.03, 29.49, 24.94, 22.99, 22.19, 14.19, 1.16.

HR-MS (ESI+): m/z $[M+H]^+$ calculated for $C_{15}H_{23}N_2O_5$: 311.1602, found: 311.1597

HR-MS (ESI-): m/z $[M-H]^-$ calculated for $C_{15}H_{21}N_2O_5$: 309.1456, found: 309.1454

### Analytical and Biochemistry methods

#### Culturing of HEK293T cells

HEK293T (CVCL_0063) cells were cultured in Dulbeccos Modified Eagles Medium – high glucose (DMEM) supplemented with 10% fetal bovine serum (FBS) and 2 mM L-alanyl-L-glutamine at 37 °C and 5% $CO_2$ atmosphere.

#### Geltrex coating of dishes for iPSCs and neuronal differentiation

Frozen Geltrex aliquots were thawn at 4 °C for several hours to O/N. Geltrex was added to cold coating medium (StableCell™ DMEM/F12) in a 1:100 dilution, mixed and immediately distributed across dishes. Coating medium volumes were 3.5 mL for p60, 10 mL for p100 and 25 mL for p150 dishes.

#### Maintenance of human iPSCs

Human iPSCs were cultured in in Geltrex coated dishes with complete StemFlex medium (StemFlex basal medium + StemFlex supplement + 1% Antibiotic-Antimycotic) at 37 °C, 5% $CO_2$. At the day of seeding or splitting, culture medium was additionally supplemented with RevitaCell (1:100), which was replaced by complete StemFlex medium the next day.

#### Differentiation of human iPSCs

$2.5*10^6$ cells were seeded in Geltrex coated p100 dishes in 10 mL complete StemFlex medium + RevitaCell (1:100) + Doxycycline (1:2000) and incubated for 24 h at 37 °C, 5% $CO_2$ (Day 1). The next day, culture medium was exchanged to 10 mL complete StemFlex and complete Neurobasal-A medium 1:1 (5 mL each) + NeuroBrew21 (1:50) + Doxycycline (1:2000). Complete Neurobasal-A medium was prepared by adding 200 mM L-Ala-L-Gln and 1% Antibiotic-Antimycotic to Neurobasal-A medium. Cells were incubated for 24 h (Day 2). The next day, culture medium was exchanged to 10 mL complete Neurobasal-A medium + NeuroBrew21 (1:50) + Doxycycline

(1:2000) (Day 3). From here medium was exchanged every second day with 10 mL fresh complete Neurobasal-A medium + NeuroBrew21 (1:50) + Doxycycline (1:2000). On Fridays, 15 mL medium was used (“double feeding”) to avoid medium change during the weekend.

#### Probe treatment *in cellulo*

Cells were either seeded with 1 * $10^6$ cells in p60 , 2 * $10^6$ cells in p100 (2.5*$10^6$ for iPSCs) or 4 * $10^6$ cells in p150 dishes in 5 mL, 10 mL or 20 mL of culture medium. Cells were either treated with pE64d (2.5 – 100 µM, varying by experiment) or respective amount of DMSO as control. After probe addition, cells were incubated 4 h prior to harvest. For the competition experiment, E64d was added 24h prior to the addition of pE64d at the concentrations mentioned in the text. Cells were harvested by washing twice with 4 or 8 mL cold DPBS (4 °C), scraped in 0.5 mL (p60) or 1 mL (p100) or 2 mL (p150) DPBS and centrifuged at 200 x g, 4 °C, 5 min. After removal of the supernatant, the cell pellets were frozen at -80 °C or directly subjected to cell lysis.

#### Cell lysis

Cells were lysed with 100 – 400 µL of lysis buffer (1% NP-40, 0.2% SDS, 20 mM Hepes, pH 7.5) via rod sonication (10 s, 20% intensity). The lysates were clarified by centrifugation (13,000 x g, 4 °C, 10 min) and supernatant was transferred into a new Eppendorf tube. The lysates were stored at -80 °C or processed directly.

#### Protein concentration measurement

Protein concentration measurement was performed with a Pierce™ BCA Protein Assay Kit (Thermo Scientific).

#### In-gel fluorescence analysis

100 µg protein per sample/replicate was diluted with lysis buffer (1% NP-40, 0.2% SDS, 20 mM Hepes, pH 7.5) ad 93.4 µL. 1 µL TAMRA-$N_3$ (10 mM in DMSO), 3 µL TCEP (100 mM in MS water) and 0.6 µL TBTA (16.7 mM in DMSO) were added and the sample was vortexed and shortly spun down. 2 µL 50 mM $CuSO_4$ was added to 100 µL final volume and the sample was vortexed, shortly spun down and incubated 1.5 h at rt, 650 rpm protected from light. Proteins were precipitated by the addition of 400 µL cold Acetone (-20 °C) and incubation at -20 °C for 1 h to O/N. The precipitated proteins were centrifuged for 10 min, 13.000 x g, 4 °C and the supernatant was removed. Proteins were reconstituted in 80 µL 0.2% SDS in PBS and sonicated 5 sec at 20% intensity with a rod sonicator. For SDS PAGE 20 µg protein per sample (16 µL) were mixed with 4 µL 5x Laemmli buffer + DTT (200 µL 5x LB + 20 µL 1M DTT) and heated for 5 min at 95 °C. Samples were loaded on a 10% self-made SDS-PAGE gel after having cooled down to rt and the gel was run for 10 min at 100V and then 50 – 60 min at 150V in 1x running buffer (25 mM Tris, 0.192 M glycine, 0.1% (m/v) SDS). In-gel fluorescence was scanned with Amersham Imager 680 (GE Healthcare). For loading control, proteins were stained with a coomassie staining solution

(0.25% Coomassie Blue R-250, 10% acetic acid, 50% MeOH) and properly destained with destaining solution (20% MeOH, 10% acetic acid).

### Western blot analysis

After running the SDS PAGE the gel was washed 3x with deionized water and the stacking gel was removed. The membrane was equilibrated in MeOH for 1 min and then the gel, blotting papers and the membrane were equilibrated in semi dry transfer buffer (5.82 g Tris, 2.9 g Glycine, 0.375 g SDS, 20% MeOH in 1 L deionized water) for 10 min. The blotting sandwich was assembled, and proteins were blotted over 30 min at 25V. After complete blotting the membrane was incubated in blocking buffer (5% milk powder in PBS + 0.5% Tween 20) for 1 h at rt, shaking. Subsequently, the membrane was incubated O/N with 10 mL of the respective primary antibody (1:1000 in blocking buffer) at 4 °C, shaking. The next day, the membrane was washed 3x with PBS + 0.5% Tween 20 for 5 min and then incubated with HRP-coupled secondary antibody (1:10.000 in blocking buffer) and anti-GAPDH rhodamine coupled antibody (1:10.000 in blocking buffer) simultaneously for 1 h at rt, protected from light. The secondary antibody and anti-GAPDH antibody were removed, and the membrane was washed 3x 10 min with PBS + 0.5% Tween 20. Chemiluminescence was scanned with Amersham Imager 680 (GE Healthcare) after activation of the membrane with Amersham ECL Prime Western Blotting Detection Reagent (1:1).

### PNGase treatment

For removing almost all *N*-linked oligosaccharides from glycoproteins, PNGaseF (NEB) was employed. Glycoprotein (20 µg), 1 µl of Glycoprotein Denaturing Buffer (10X), and $H_2O$ were combined to make a 10 µl total reaction volume. Glycoproteins were denatured by heating the reaction at 100 °C for 10 min. Denatured samples were spun down, and 2 µl GlycoBuffer 2 (10×), 2 µl 10% NP-40, and 6 µl $H_2O$ were added to make a total reaction volume of 20 µl. Then, 1 µl PNGase F was added, the reaction mixture was gently mixed and incubated for 1 h at 37 °C, and successful deglycosylation was assessed by SDS-PAGE and subsequent WB analysis.

### Polymerase Chain Reaction (PCR)

PCR was performed for cloning purposes using plasmid DNA (pcDNA3.1_hCTSB, RRID: Addgene_11249) as a template. Here, the procedure was performed according to the manufacturers' protocol (Q5 site–directed mutagenesis, NEB #E0554S). The thermocycling conditions applied are listed in Table S3. PCR product formation was validated by 0.7 % agarose gel electrophoresis.

### 0.7% Agarose Gel

For validation of PCR product formation, 1% agarose gel electrophoresis was conducted. The agarose was dissolved in 40 ml 1× 40 mM Tris, 20 mM acetic acid, 1 mM EDTA buffer, and heated up in the microwave. After cooling down, 2 µl Gel stain was added and the gel was polymerized. As a standard, 1 kB DNA ladder (Carl Roth) was used. To the PCR product, 6× stain

(NEB) was added (cf = 1×). The electrophoresis was conducted for 1 h at 100 V in 0.5× Tris, acetic acid, and EDTA buffer. PCR products were detected under UV light.

#### KLD Reaction

A KLD (Kinase, Ligase, DpnI) reaction was performed with the PCR product according to the manufacturers' protocol (Q5 site–directed mutagenesis, NEB #E0554S). Here, the PCR product is first phosphorylated (kinase), then ligated (ligase), and residual template DNA is decomposed by DpnI enzyme (only methylated DNA is a substrate). The reaction mixture was prepared, mixed by pipetting up and down 5 to 10 times, and incubated for 5 min at room temperature. The KLD mixture was used for subsequent bacterial transformation.

#### Chemical Transformation of Bacterial Cells

For chemical transformation, chemically competent E. coli DH5-alpha cells stored at −80 °C were thawn for 10 min on ice. Then, 5 µl of the Kinase, Ligase, and DpnI (KLD) reaction mixture was added directly to 50 µl of cells, and the tube was gently flicked three times. The cells were incubated for 30 min on ice and then heat shocked at 42 °C for 30 s. shock. The cells were immediately placed on ice for 5 min for regeneration before the addition of 950 µl of SOC medium (NEB) and incubation for 1 h at 37 °C at 250 rpm. In the meantime, LB-Agar (1.3% Agar) plates supplemented with 100 µg/ml ampicillin were prewarmed at 37 °C. After incubation, 100 µl of the cell suspension was spread onto prewarmed LB-Agar plates and evenly distributed using glass beads by shaking the plate under sterile conditions. The LB-Agar plates were then incubated overnight at 37 °C and subsequently stored at 4 °C or directly used for plasmid amplification.

#### Q5 Site–Directed Mutagenesis

For preparation of hCTSB C108A mutant starting from pcDNA3.1_hCTSB (RRID:Addgene_11249), the Q5 Site-Directed Mutagenesis Kit (New England Biolabs #E0554S) was used. The sequence was validated by Sanger sequencing (Genewiz) using the sequencing oligonucleotide listed in Table S4. Sequences of primers used for the introduction of the C108A mutation are shown in Table S5.

#### Plasmid DNA Isolation Using Mini/Midi Preparation Kit

For plasmid DNA isolation, either the Mini (small scale, NEB) or Midi (large scale, Zymo) preparation kit was utilized. For Mini or Midi preparation, 3 ml or 100 ml, respectively, of sterile LB medium supplemented with 100 µg/ml ampicillin was prepared. A single clone from the overnight LB agar plate was selected using a pipette tip and subsequently added to the medium. The medium was then incubated for 12 to 16 h at 37 °C at 180 rpm. Plasmid DNA was isolated following the manufacturer's protocol for the Plasmid Miniprep/Midiprep Kit. The DNA was eluted in the respective elution buffer included in the kit. DNA concentration was measured using a Nanodrop, and the DNA sequence was confirmed by Sanger sequencing conducted by Genewiz.

**Enrichment of modified proteins using the SP2E protocol for LC-MS/MS**

400 µg protein per sample/replicate was diluted with lysis buffer (1% NP-40, 0.2% SDS, 20 mM Hepes, pH 7.5) add 186.8 µL. 2 µL Biotin-$N_3$ (10 mM in DMSO), 6 µL TCEP (100 mM in MS water) and 1.2 µL TBTA (16.7 mM in DMSO) were added and the sample was vortexed and shortly spun down. 4 µL 50 mM $CuSO_4$ was added to 200 µL final volume and the sample was vortexed, shortly spun down and incubated for 1.5 h at r.t., 650 rpm. After the incubation was completed, 200 µL of 8 M urea in MS-water was added to the click reaction to a final volume of 400 µL. 100 µL of a 1:1 mixture of hydrophobic and hydrophilic carboxylate-coated magnetic beads (Cytiva) was washed thrice with 500 µL of MS-water and the reaction mixture was transferred onto the pre-washed beads followed by the addition of 600 µL absolute Ethanol, mixing and incubation at r.t., 950 rpm for 5 min. The beads were washed thrice with 500 µL of 80% EtOH in MS-water. Subsequently, proteins were eluted from the carboxylate-coated beads twice with 500 µL of 0.2% SDS in PBS at r.t., 950 rpm for 5 min and transferred onto 50 µL streptavidin-coated magnetic beads (New England Biolabs) that had been equilibrated 3x with 500 µL 0.2% SDS in PBS. For streptavidin-biotin complex formation, beads were incubated at r.t., 950 rpm for 1 h with eluted proteins. After the incubation, beads were washed three times with 500 µL of 0.1% NP-40 in PBS, two times with 500 µL of 6 M urea (in MS-water) and two times with 500 µL of MS-water by vortexing and short-spin between each wash step. After the last wash step, 80 µL of ABC buffer (ammoniumbicarbonate, 125 mM in MS-water), 10 µL of TCEP (100 mM in MS-water) and 10 µL of chloroacetamide (400 mM in MS-water) was added and incubated at 95 °C for 5 min. Then, proteins were digested O/N with 1.5 µL sequencing-grade trypsin (0.5 mg/mL, Promega) at 37 °C, 650 rpm. The next day supernatants were transferred into new Eppendorf tubes, and beads were washed thrice with 100 µL ABC buffer (100 mM in MS-water) to collect the peptides (wash fractions were combined into the corresponding tubes). The combined fractions were acidified with 2 µL of MS-grade FA and peptides were desalted with 50 mg of SepPak C18 cartridges on a vacuum manifold. First, cartridges were equilibrated with 1 mL of MS grade ACN, 1 mL of elution buffer (80% ACN with 0.5% FA in MS-water) and 3 mL of wash buffer (0.5% FA in MS-water). Second, combined peptide fractions were loaded onto the cartridges and washed with 3 mL of wash buffer. Finally, peptides were eluted twice with 250 µL of elution buffer. Desalted peptide eluates were vacuum-dried with a SpeedVac at 35 °C, reconstituted in 30 µL of 1% FA in MS-water (vortex and 15 min sonication bath), transferred to MS vials and subjected to LC-MS/MS analysis with 2 – 5 µl injection volume.

**Enrichment of modified proteins using the SP2E protocol for SDS PAGE & Western Blot**

300 µg protein per sample/replicate was diluted with lysis buffer (1% NP-40, 0.2% SDS, 20 mM Hepes, pH 7.5) add 93.4 µL. 1 µL TAMRA-Biotin-$N_3$ (10 mM in DMSO), 3 µL TCEP (100 mM in MS water) and 0.6 µL TBTA (16.7 mM in DMSO) were added and the sample was vortexed and shortly spun down. 2 µL 50 mM $CuSO_4$ was added to 100 µL final volume and the sample was vortexed, shortly spun down and incubated for 1.5 h at r.t., 650 rpm, protected from light. After the incubation was completed, 100 µL of 8 M urea in MS-water was added to the click reaction to a final volume of 200 µL. 100 µL of a 1:1 mixture of hydrophobic and hydrophilic carboxylate-coated magnetic beads (Cytiva) was washed thrice with 500 µL of MS-water and the reaction mixture was transferred onto the pre-washed beads followed by the addition of 600 µL absolute Ethanol, mixing and incubation at r.t., 950 rpm for 5 min. The beads were washed thrice with 500 µL of 80% EtOH in MS-water. Subsequently, proteins were eluted from the carboxylate-coated beads twice with

500 µL of 0.2% SDS in PBS at r.t., 950 rpm for 5 min and transferred onto 50 µL streptavidin-coated magnetic beads (New England Biolabs) that had been equilibrated 3x with 500 µL 0.2% SDS in PBS. For streptavidin-biotin complex formation, beads were incubated at r.t., 950 rpm for 1 h with eluted proteins. After the incubation, beads were washed three times with 500 µL of 0.1% NP-40 in PBS, two times with 500 µL of 6 M urea in MS-water and two times with 500 µL of MS-water by vortexing and short-spin between each wash step. After the last wash step, the beads were resuspended in 50 µL 1x Laemmli Buffer supplemented with 20 mM DTT and incubated for 5 min at 95 °C. The beads were removed on a magnetic rack, and the supernatant was transferred into a new Eppendorf tube. For unprocessed lysates 20 µg protein was diluted in 0.2% SDS in PBS ad 16 µl and 4 µL 5x Laemmli Buffer + 100 mM DTT was added (final 1x LB + 20 mM DTT) and incubated for 5 min at 95 °C. 20 µL of enriched or unprocessed sample was loaded onto a 10 % SDS gel. The PAGE was run at 100V for 10 min, and then at 150V for ~60 min, while protected from light. Subsequently, Western blot analysis was performed as previously described.

**Enrichment of modified peptides using desthiobiotin-tag**

Probe or DMSO treated cells were harvested by washing twice with 8 mL cold DPBS (4 °C), scraped in 3 mL cold DPBS and then 3x 1 mL of this suspension was split in Eppendorf tubes, yielding 3 technical replicates of each probe treated and DMSO treated samples. Each replicate was lysed in 1300 µL lysis buffer (1% NP-40, 0.2% SDS, 20 mM Hepes, pH 7.5) as previously described and protein concentration was determined by Pierce™ BCA Protein Assay Kit (Thermo Scientific). A total amount of 8 - 10 mg protein of each control and *in cellulo* probe-treated lysates (split across three replicates each) was used for the subsequent workflow. All replicates were diluted with lysis buffer ad 1893.5 µL. 4 µL Desthiobiotin-$N_3$ (50 mM in DMSO), 60 µL TCEP (100 mM in MS water) and 2.5 µL TBTA (83.5 mM in DMSO) were added, and the sample was vortexed and shortly spun down. 40 µL 50 mM $CuSO_4$ was added to 2 mL final volume, and the sample was vortexed, shortly spun down and incubated 2.5 h at rt, 650 rpm. Subsequently, the click reaction mixture was transferred into a 15 ml falcon tube, 8 mL cold acetone (-20 °C) was added and proteins were precipitated at -20 °C O/N. Precipitated proteins were spun down 10 min, 13,000 x g, 4 °C, supernatant was disposed, and each protein pellet was washed twice with 1 mL of cold methanol by resuspending the pellet with sonication (3x, 5 s, 20% intensity) and centrifugation. After the last wash step, methanol was removed, and the pellet was air-dried for 10 min. The pellet was reconstituted in 300 µL of 8 M urea in 0.1 M TEAB solution by sonication (3x, 10 s, 20% intensity) and diluted with 900 µL of 0.1 M TEAB ad 1.2 mL total volume. 100 µL of streptavidin-coated agarose beads (Sigma Aldrich) per sample (probe and control) were pre-washed thrice with 1 mL of 0.2% NP-40 in PBS. The washed beads were split equally across 2x three Eppendorf tubes (3x probe, 3x control). The reconstituted replicates were added to the beads and incubated for 1.5 h at rt, 850 rpm. All the subsequent washing steps were performed by shortly vortexing the tubes, centrifuging 2 min, 2000 rpm, r. t. and disposal of the supernatant. For enrichment, the beads were washed thrice with 1 mL 0.1% NP-40 in PBS (the three replicates of each sample were combined after the first washing step), thrice with 1 mL of PBS, thrice with 1 mL of MS-grade $H_2O$ and once with 1 mL 2 M urea in 100 mM ABC buffer. After the last supernatant was discarded, the beads were resuspended in 190 µL 2 M urea in 100 mM ABC buffer, 2 µL of 1 M DTT was added and incubated at 37 °C for 45 min, 1000 rpm. The reduced thiol groups were alkylated by the addition of 8 µL 0.5 M iodoacetamide (fresh prepared with 2 M urea in 100 mM ABC buffer) and incubation at r. t. for 1 h. 600 µL 100 mM ABC buffer was added, vortexed, centrifuged at 1000 x g, 2 min and the supernatant was removed. For digestion, beads were resuspended in 100 µL 100 mM ABC buffer, then 4 µL of sequencing-grade trypsin (0.5 mg/mL, Promega) was added and proteins were digested at 37 °C, 1000 rpm, O/N. The next day, 800 µL 0.1% NP-40 in PBS was added to the tryptic digest, centrifuged and supernatant was removed. The beads were washed thrice with 1 mL of 0.1% of NP-40 in PBS, thrice with 1 mL PBS and thrice with 1 mL MS grade $H_2O$. Desthiobiotin-modified peptides were eluted twice with 200 µL of ACN/$H_2O$ (1:1) with 0.1% FA, twice with 100 µL of ACN/$H_2O$ (1:1) with 0.5% FA and twice with 100 µL of ACN/$H_2O$ (4:1) with 0.5% FA. For each step, beads were incubated with the corresponding elution buffer for 10 min, 1000 rpm, 40 °C, spun down and the supernatant eluate fraction collected and combined. Combined fractions were centrifuged for 1 min, 20,000 x g and transferred into a new Eppendorf tube while ~50 µL of the eluate was left to ensure that no beads were collected. The eluate was vacuum-dried with a SpeedVac at 35 °C, reconstituted in 20 µL of 1% FA in $H_2O$, vortexed and sonicated for 15 min. Finally, peptide mixtures were transferred to MS-vials and submitted to LC-MS/MS measurement with an injection volume of 8 µL.

**Whole proteome analysis using the SP3 protocol**

10 µg protein per replicate was diluted with lysis buffer (1% NP-40, 0.2% SDS, 20 mM Hepes, pH 7.5) add 10 µL in a 96 well plate. A 1:1 mixture of hydrophobic and hydrophilic carboxylate-coated magnetic beads (Cytiva) was washed thrice with 500 µL of MS-water and resuspended to the initial volume. 2 µl of these pre-washed beads were added to the proteins, followed by the addition of 48 µL absolute Ethanol and 10 min incubation at rt, 1000 rpm. The following washing steps were performed by the Microlab prep pipetting robot (Hamilton). The 96 well plate was transferred to a magnetic rack and incubated for 2 min. The supernatant was discarded. Beads were washed twice with 180 µL 80% EtOH (30s incubation, rt, shaking). Beads were washed once with 180 µL of ACN (LC-MS grade, 30s incubation, rt, shaking). Supernatant was discarded and beads were air-dried for 30s. The beads were reconstituted in 20 µL 100 mM ABC buffer and 1 µL trypsin (0.5 mg/mL, Promega) was added and proteins were digested at 37 °C, 1000 rpm, O/N. The next day beads were resuspended by pipetting, and the supernatant was removed on a magnetic rack and transferred into new Eppendorf tubes. Proteins were eluted twice by the addition of 10 µL MS-water and 5 min incubation at 40 °C. 4 µL 1% formic acid in MS water was added to the combined fractions, tubes were placed on a magnetic rack and peptides were transferred into MS vials while ~ 20 µL were left to ensure that no beads were collected. Samples were subjected to LC-MS/MS analysis with 1 µL injection volume.

**IRAK1 immunoprecipitation**

6 * $10^6$ HEK293T cells were seeded in p150 dishes with 20 mL of culture medium and cultured until ~ 90% confluency. Cells were then treated with 50 µM pE64d or 50 µM E64d (10 µL of 100 mM stock), or 10 µL DMSO as control. After the treatment, cells were incubated for 4 h and then harvested by removing the culture medium, washing twice with 8 mL cold (4 °C) DPBS, addition of 3 mL ice cold (- 20 °C) fresh prepared IP-lysis buffer (1% NP-40, 50 mM Tris/HCl, pH 7.4, 150 mM NaCl, 1x cOmplete Protease Inhibitor), and scraping from the dish. The cell suspension was equally distributed across three Eppendorf tubes and stored on ice to yield 3 replicates per sample (3 x 3 = 9 replicates overall). The cells were then lysed, and the protein concentration was

determined as previously described. For protein-antibody-complex formation, 3 mg protein per replicate (9 mg protein overall per sample) was diluted with IP-lysis buffer ad 1.3 mL and 4 µg anti-IRAK1 antibody (Proteintech 10478-2-AP) was added to each replicate. This mixture was incubated O/N at 4 °C with gentle agitation. The next day, 25 µL/replicate (0.25 mg) Pierce™ Protein A/G magnetic beads (10 mg/ml) were transferred into an Eppendorf tube, and 175 µL ice-cold IP-lysis buffer was added, resuspended by pipetting and the supernatant was removed on a magnetic rack. The beads were washed three times with 1 mL ice-cold IP-lysis buffer by vortexing, but not short spinning to prevent aggregation of the beads and potential loss of binding affinity. The protein-antibody mixture was added to the beads, vortexed, and incubated at 4 °C for 4 h with gentle agitation. The three replicates of each sample were combined, and the supernatant was removed on a magnetic rack. Subsequently, the beads were washed three times by vortexing and short spinning with 1 mL ice-cold IP-lysis buffer, followed by washing three times with 1 mL ice-cold IP-wash buffer (50 mM Tris/HCl, pH 7.4, 150 mM NaCl) and once with MS water. For the digestion of enriched proteins, beads were resuspended in 100 µL of 100 mM ABC buffer supplemented with 4 µL of trypsin (0.5 µg/µL, Promega) and incubated overnight at 37 °C with shaking at 650 rpm. Digested peptides were transferred to new Eppendorf tubes and beads were washed three times with 100 µL of 100 mM ABC buffer. The combined elute fractions were acidified with 2 µL formic acid (FA) and subsequently desalted on a vacuum manifold. Therefore, SepPak C18 Cartridges (50 mg columns, Waters #186000308) were equilibrated with 1 mL ACN, then 1 mL of 80% ACN containing 0.5% FA, and then 3x with 1 mL of 0.5% FA in MS-grade water. Samples were loaded without vacuum, washed three times with 1 mL of 0.5% FA in MS-grade water under vacuum, and eluted twice with 250 µL of 80% ACN containing 0.5% FA without vacuum. The solvent was removed with SpeedVac at 37 °C and peptides were dissolved in 30 µL of 1% FA in MS-grade water, followed by sonication for 15 min. Finally, the samples were transferred into MS-vials.

**LC-MS/MS measurement**

MS measurements were performed on a Orbitrap Eclipse Tribrid Mass Spectrometer (Thermo Fisher Scientific) coupled to an UltiMate 3000 Nano-HPLC (Thermo Fisher Scientific) via a Nanospray Flex (Thermo Fisher Scientific) and FAIMS interface (Thermo Fisher Scientific). First, peptides were loaded on an Acclaim PepMap 100 µ-precolumn cartridge (5 µm, 100 Å; 300 µM ID x 5 mm, Thermo Fisher Scientific). Then, peptides were separated at 40 °C on a PicoTip emitter (noncoated, 15 cm, 75 µm ID, 8 µm tip, New Objective) that was in house packed with ReprosilPur 120 C18-AQ material (1.9 µm, 150 Å, Dr. A. Maisch GmbH) or on an Aurora Elite XT C18 UHPLC column (15 cm, 75 µm ID, IonOpticks). The LC buffers consisted of MS-grade water (A) and acetonitrile (B) both supplemented with 0.1% formic acid. The gradient was run from 4-35.2% B during a 60 min method (0-5 min 4%, 5-6 min to 7%, 7-36 min to 24.8%, 36-41 min to 35.2%, 40-41.1 min 80%, 41.1-46 min 80%, 46-60 min 4%) at a flow rate of 300 nL/min.

*Data-independent acquisition for SP2E enrichment*

FAIMS was performed with one CV at -45 V. One DIA cycle comprised one MS1 scan followed by 30 MS2 scans. The mass spectrometer was operated in DIA mode with following settings: Polarity: positive; MS1 Orbitrap resolution: 60k; MS1 AGC target: standard; MS1 maximum injection time: 50 ms; MS1 scan range: m/z 200-1800; RF Lens: 30%; Precursor Mass Range: m/z 500-740; isolation window: m/z 4; window overlap: m/z 2; MS2 Orbitrap resolution: 30k; MS2 AGC target:

200%; MS2 maximum injection time: auto; HCD collision energy: 35%; RF Lens: 30%; MS2 scan range: auto.

*Data-dependent acquisition for Desthiobiotin enrichment (Site-ID)*

The mass spectrometer was operated without FAIMS in DDA mode with following settings for MS1: Polarity: positive; Orbitrap Resolution: 240k; Mass Range: Normal; Scan Range: 375 – 1500; RF Lens: 30%; AGC Target: Standard; Maximum Injection Time Mode: Auto; Microscans: 1; Intensity Threshold: 1.0e4; Included charge state(s): 2-6; Dynamic Exclusion: 30s; Mass Tolerance: 10 ppm. Top 10 precursors were selected for fragmentation in MS2 with following settings: Orbitrap Resolution: 15k; Mass Range: Normal; First Mass (m/z): 110; AGC Target: 200%; Maximum Injection Time Mode: Auto; HCD Collision Energy: 30%. A Targeted Mass Trigger Filter was applied for m/z 197.1284 (Dehydrodesthiobiotin, D1) and m/z 240.1706 (DTB-Oxonium Ion, D2) with the following settings: Mass Tolerance: 25 ppm; Number of Scans: 2, Orbitrap Resolution: 60k; Mass Range: Normal; First Mass (m/z): 110; AGC Target: 200%; Maximum Injection Time Mode: Auto; HCD Collision Energy: 30%.

*Data-independent acquisition for IRAK1 immunoprecipitation (Site-ID)*

The mass spectrometer was operated without FAIMS in DIA mode with following settings for MS1: Polarity: positive; Orbitrap Resolution: 120k; Mass Range: Normal; Scan Range: 375 – 1500; RF Lens: 30%; AGC Target: Standard; Maximum Injection Time Mode: Auto; Microscans: 1 and for MS2: Orbitrap Resolution: 30k; Precursor Mass Range (m/z): 500 - 740; First Mass (m/z): 110; RF Lens: 30%; AGC Target: 200%; Maximum Injection Time Mode: Auto; Microscans: 1; HCD Collision Energy: 30%, Number of Spectra: 30.

**Computational evaluation of measured mass spectra**

DIA-NN (peptide quantification)

Measured .raw files were analysed with DIA-NN 2.1 and peptides were searched against Uniprot database for Homo sapiens (UP000005640, taxon identifier: 9606) with included isoforms The DIA-NN[2] settings were as follows: Contaminants: enabled; FASTA digest for library-free search/library generation: enabled; Deep learning-based spectra, RTs and IMs prediction: enabled; missed cleavages: 2; max number of variable modifications: 2; modifications: N-term M excision, carbamidomethylation, oxidation (M) and N-term acetylation; Peptide length range: 7 – 35; Precursor charge range: 2-6; precursor range: m/z 500-740; fragment ion range: m/z 200-1800; precursor FDR level: 1%; match between runs (MBR): enabled; all other parameters were used as default.

FragPipe platform (Site-ID)

MS *.raw files were converted to *.mzML format with “MSConvert” with following settings: “peakPicking” filter with “vendor msLevel = 1”, “titleMaker” filter was enabled. Peptides were searched in Fragpipe 23.1 against Uniprot database for Homo sapiens (UP000005640, taxon identifier: 9606) with included contaminants and decoys. Due to the huge variety of settings, the four general FragPipe workflows (open, closed, offset search and diagnostic ion mining) are provided on the PRIDE server. In the following most important features were listed:

*Open search*: Precursor mass tolerance: -150 - 800 Da, variable modifications: carbamidomethylation (C), oxidation (M) and acetylation (N-term); fixed modifications: all enabled with Mass Delta 0

*Closed search:* variable modifications: 696.3806 (C), carbamidomethylation (C), oxidation (M) and acetylation (N-term); fixed modifications: none

*Offset search*: variable modifications: 696.3806 (C), carbamidomethylation (C), oxidation (M) and acetylation (N-term); fixed modifications: all enabled with Mass Delta 0; mass offset: 0/396.3806

*Diagnostic ion mining*: Precursor mass tolerance: -0 - 700 Da, variable modifications: none; fixed modifications: enabled on all sites except C with Mass Delta 0; Run PTM-Sheperd: enabled; Mine for diagnostic ions and fragments: enabled; Extract known diagnostic ions from spectra: enabled; Diagnostic fragment masses: 197.12845, 240.17065, 583.39261, 731.37546

**Data statistics and evaluation from computational output**

Statistics and plots were generated using a custom R script. In brief, the “report.parquet” file obtained from the DIA-NN calculation was loaded as the main data using the readDIANN function from the limpa package[3] with q.cutoffs set to 0.01. Next, non-proteotypic peptides were filtered out and groups were added to specify control or probe treated samples. The detection-probability curve was generated using the dpc function and proteins were quantified from the “Protein.Names” column using dpcQuant (for heatmaps proteins were quantified from the “Genes” column to avoid multiple entries for a single gene)[3]. Differential expression analysis was performed using the limma package[4]. Volcano plots were generated using the EnhancedVolcano package and cutoffs were set to 0.05 for the adjusted p-value (corrected using the Benjamin Hochberg method) and 0.01 for log2 fold change. Heatmaps were generated using the pheatmap package. Boxplots and PCA plots were generated using the ggplot2 package. Profile plots of individual peptides were generated using the plotPeptide function from the limpa package.

*Open search for modified peptides (FragPipe)*: Mass shifts of modified peptides were obtained from the “psm.tsv” file “observed modifications” column.

*Closed search for modified peptides (FragPipe)*: Mass shifts of modified peptides were obtained from the “psm.tsv” file “assigned modifications” column.

*Offset search for modified peptides (FragPipe)*: Mass shifts of modified peptides were obtained from the “psm.tsv” file “assigned modifications” column.

*Diagnostic ion mining for modified peptides and diagnostic ions (FragPipe)*: Diagnostic ion masses were obtained from the “global.diagmine” file “mass” column and were drawn in ChemDraw to match the obtained masses to the ion structure. The number of PSMs for the intact modification and the diagnostic ions in sample P_wt were extracted from the “P_wt.diagnosticProfile” file (see also Table S1)

**Materials**

| Reagent/Resource | Reference or Source | Identifier or Catalog Number |
|---|---|---|
| **Experimental Models** | | |
| Human: HAP-1 | Prof. Dr. Lucas Jae | |
| Human: HEK293T | Ref.[5] | RRID: CVCL_0063 |
| Human: iNGNs | Prof. Dr. Thomas Carell | |
| **Recombinant DNA** | | |
| pcDNA3.1_hCTSB | Hyeryun Choe Lab | RRID:Addgene_11249 |
| pcDNA3.1_hCTSB_C108A | This study | Derived from Addgene_11249 |
| **Antibodies** | | |
| Anti-CTSB | proteintech | Cat# 12216-1-AP |
| Anti-PLD3 | Sigma-Aldrich | Cat# HPA012800 |
| Anti-BLMH | proteintech | Cat# 14941-1-AP |
| Anti-IRAK1 | Cell Signaling | Cat# 4359S |
| Anti-PSMA1 | proteintech | Cat# 11175-1-AP |
| Anti-SUPT5H | proteintech | Cat# 16511-1-AP |
| Anti-CTSZ | proteintech | Cat# 16511-1-AP |
| Anti-GAPDH-Rhodamine | Bio-Rad | Cat# 12004168 |
| Goat-anti-rabbit HRP | Thermo Fisher | Cat# 31460 |
| **Oligonucleotides and other sequence-based reagents** | | |
| / | | |

| **Chemicals and Enzymes** | | |
|---|---|---|
| Acetic acid (100%) | | |
| Acetone (HPLC grade) | VWR Chemicals | Cat# 20067.320 |
| Acetonitrile (LC-MS grade) | Thermo Fisher | Cat# A955-212 |
| Agar-Agar | Carl Roth | Cat# 6494.3 |
| Alanyl-Glutamine | Sigma-Aldrich | Cat# G8541 |
| Ammonium Acetate | Sigma-Aldrich | Cat# 09689-1kg |
| Ammonium bicarbonate | Fluka | Cat# 09830-100g |
| Ammoniumperoxodisulfat | Sigma-Aldrich | Cat# 09913 |
| Ampicillin | Carl Roth | Cat# HP62.1 |
| Antibiotic-Antimycotic | Gibco | Cat# 15240062 |
| Avidin-agarose beads | Sigma-Aldrich | Cat# A9207-5ML |
| Benchmark™ fluorescence Protein Standard | Invitrogen | Cat# LC5928 |
| Biotin-$PEG_3$-$N_3$ | Carbosynth | Cat# FA34890 |
| BSA | AppliChem | Cat# A6588 |
| Carboxylate-coated magnetic beads (hydrophobic) | Cytiva | Cat# 65152105050250 |

| | | |
|---|---|---|
| Carboxylate-coated magnetic beads (hydrophilic) | Cytiva | Cat# 45152105050250 |
| Chloroacetamide | Sigma-Aldrich | Cat# C0267-100g |
| Color Prestained Protein Standard (10-250 kDa) | New England Biolabs | Cat# P7719S |
| cOmplete™ Protease Inhibitor Cocktail | Roche | Cat# 04693116001 |
| Coomassie Blue R-250 | Fluka | Cat# 27816 |
| $CuSO_4$ x 5 H2O | Acros | Cat# 10627162 |
| dd$H_2O$ (LC-MS grade) | Honeywell | Cat# 15665350 |
| Desthiobiotin-$PEG_3$-$N_3$ | Sigma-Aldrich | Cat# 902020-25mg |
| DIPEA | Sigma-Aldrich | Cat# 387649 |
| DMEM (1x) | Sigma-Aldrich | Cat# D6546 |
| DMF anhydrous | Sigma-Aldrich | Cat# 227056 |
| DMSO | Sigma-Aldrich | Cat# D4540 |
| Doxycyclin-hyclat | Sigma-Aldrich | Cat# D9891 |
| DPBS (1x) | Sigma-Aldrich | Cat# D8357 |
| DTT | AppliChem | Cat# A2948,0025 |
| EDTA | BioChemica | Cat# A1103 |
| Ethanol (EtOH) | Merck | Cat# 34852 |
| Fetal bovine serum | Thermo Fisher | Cat# A3840001 |
| Fetal bovine serum (FBS) | Biochrom | Cat# S0115 |
| Formic Acid (LC-MS grade) | Thermo Fisher | Cat# A117 |
| GelStain | Carl Roth | Cat# 3865.1 |
| Geltrex™ Flex LDEV-Free Reduced Growth Factor Basement Membrane Matrix | Gibco | Cat# A4000046703 |
| Glycerol | Sigma-Aldrich | Cat# G5516 |
| HCl | In house chemical store | N/A |
| Hepes | Carl Roth | Cat# HN77.5 |
| HEPES solution | Sigma-Aldrich | Cat# H0887-100ML |
| Iodoacetamide (IAA) | Sigma-Aldrich | Cat# I6125-10g |
| LB medium | Carl Roth | Cat# 66693 |
| Methanol (LC-MS grade) | Thermo Fisher | Cat# A456 |
| NaOH | Sigma-Aldrich | Cat# S5881 |
| Neurobasal-A Medium | Gibco | Cat# 10888022 |
| MACS® NeuroBrew®-21 | Mytenyi Biotec | Cat# 130-093-566 |
| NP-40 | Sigma-Aldrich | Cat# 74385 |
| Opti-MEM™ I Reduced Serum Medium, GlutaMAX™ Supplement | Thermo Fisher | Cat# 51985-026 |
| pE64d | See Organic Synthesis | N/A |
| Penicillin/Streptomycin | Gibco | Cat# 15-140-122 |
| Polyethylenimine (PEI) | Polyscience | Cat# 23966 |
| Propargylamine probe | Propargylamine probe | Propargylamine probe |

| | | |
|---|---|---|
| RevitaCell™ Supplement (100X) | Gibco | A2644501 |
| Rotiphorese.Gel 30 (37,5:1) | Carl Roth | Cat# 3029.1 |
| sodium dodecyl sulfate | AppliChem | Cat# A2572 |
| StableCell™ DMEM/F12 | Sigma-Aldrich | Cat# D0697 |
| StemFlex Medium + Supplement | Gibco | Cat# A3349401 |
| Streptavidin magnetic beads | New England Biolabs | Cat# S1420S |
| Synth-a-Freeze™ Cryopreservation Medium | Gibco | Cat# A1254201 |
| TAMRA-$N_3$ | Baseclick | Cat# BCFA-008-1 |
| TBTA | TCI | Cat# T2993 |
| TCEP | Carbosynth | Cat# FT01756 |
| TEAB (1 M) | Sigma-Aldrich | Cat# T7408 |
| TEMED | Sigma-Aldrich | Cat# T9281 |
| Tris-base | Thermo Fisher | Cat# 10724344 |
| Triton X-100 | Sigma-Aldrich | Cat# X100 |
| Trypan Blue | Thermo Fisher | Cat# 11538886 |
| TrypLE Express | Thermo Fisher | Cat# 12604013 |
| Trypsin (sequencing-grade) | Promega | Cat# V5113 |
| Tween 20 | Sigma-Aldrich | Cat# P6585 |
| Urea | AppliChem | Cat# A1049 |
| **Software** | | |
| FragPipe | Ref.[6–11] | https://fragpipe.nesvila b.org/ |
| DIA-NN | Ref.[2] | https://github.com/vde michev/DiaNN |
| MSConvert | Ref.[12] | https://proteowizard.so urceforge.io/download. html |
| PDV viewer | Ref.[13] | https://github.com/wen bostar/PDV |
| FreeStyle | Thermo Fisher Scientific | N/A |
| R | N/A | https://www.R-project.org/>. |
| RStudio | N/A | http://www.posit.co/ |
| Venny | N/A | https://bioinfogp.cnb.c sic.es/tools/venny/ |
| **Kits and Other** | | |
| Aurora Elite XT C18 UHPLC column | ionopticks | Cat# AUR4-15075C18-XT |
| Pierce® BCA Protein Assay Kit | Thermo Fisher Scientific | Cat# 23225 |
| ZymoPURE™ II Plasmid Midiprep Kit | Zymo Research | Cat# D4200 |
| Plasmid Miniprep Kit | New England Biolabs | Cat# T1010S |
| Q5 Site-Directed Mutagenesis Kit | New England Biolabs | Cat# E0554S |

| | | |
|---|---|---|
| Orbitrap Eclipse Tribrid Mass Spectrometer | Thermo Fisher Scientific | N/A |
| FAIMS Pro Duo Interface | Thermo Fisher Scientific | N/A |
| PicoTip™ Emitter, Silica Tip™ | New Objectives | FS360-75-8-N-20-C15 |
| ReproSil-Pur 120 C18-AQ, 1.9 µm | Dr. Maisch GmbH | Cat# r119.aq.0001 |
| ReproSil-Pur 120 NH2, 3 µm | Dr. Maisch GmbH | Cat# r13.a0.0001 |
| PEPMAP100 C18 5UM 0.3X5MM | Thermo Fisher Scientific | Cat# 160454 |
| Sep-Pak C18 1 cc Vac Cartridge, 50 mg | Waters | 186000308 |
| VP 250/10 Nucleodur 100-5 C18 | AppliChem | Cat# 762022.100RC |

**NMR Spectra**

Diethyl (2S,3S)-oxirane-2,3-dicarboxylate

$^1$H NMR (500 MHz, $CDCl_3$)

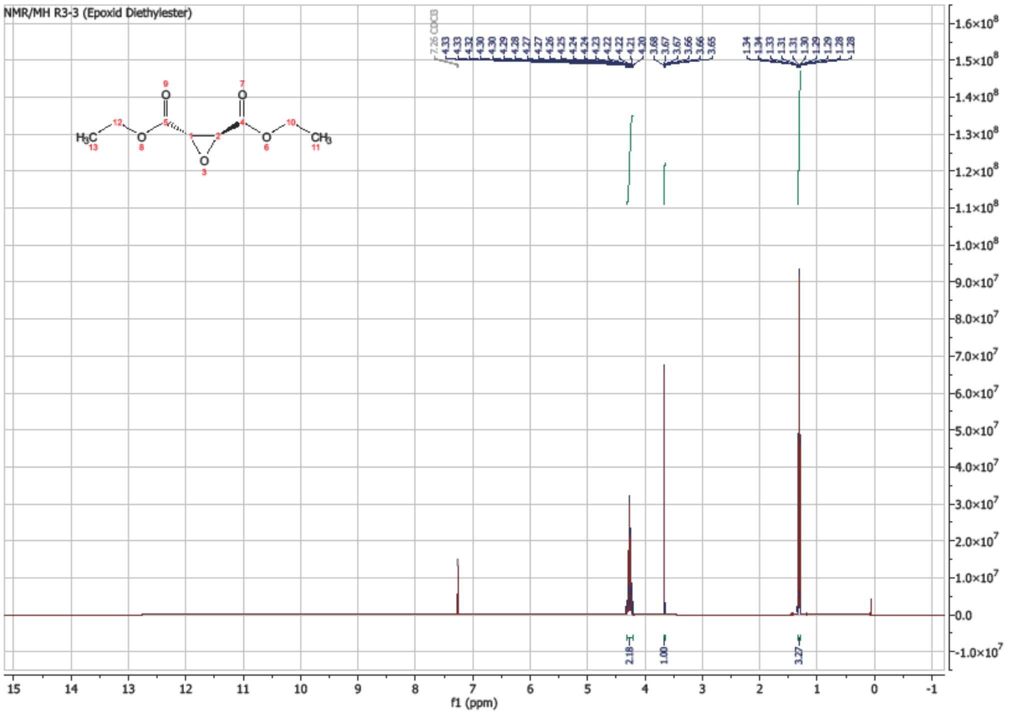

$^{13}$C NMR {$^1$H} (126 MHz, $CDCl_3$)

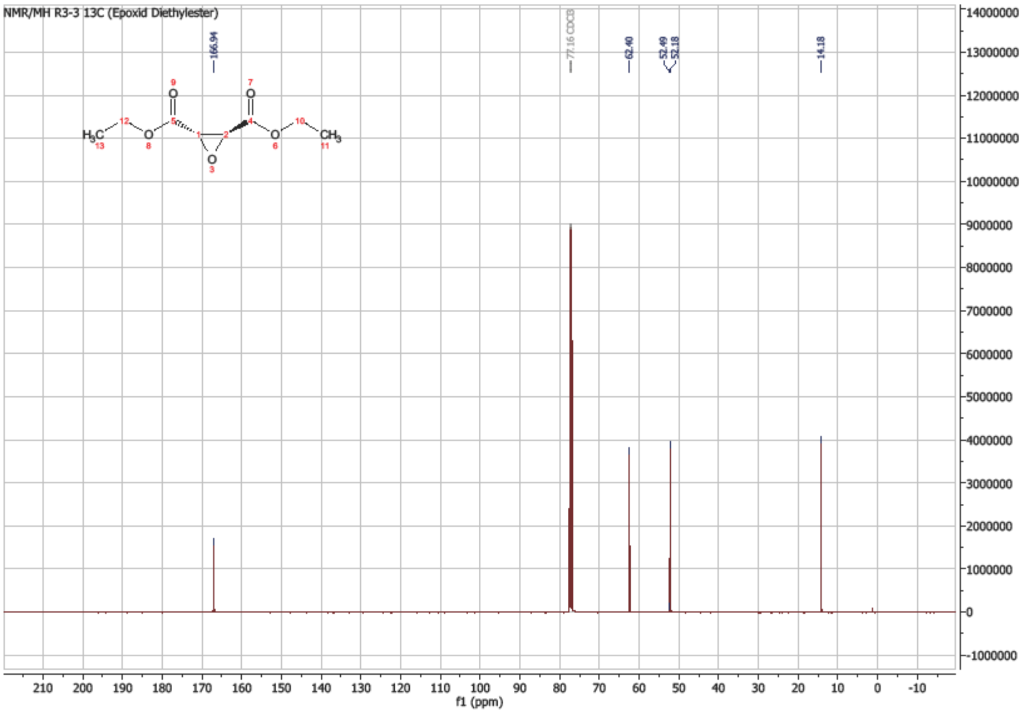

Ethyl (2S,3S)-3-((4-methyl-1-oxo-1-(prop-2-yn-1-ylamino)pentan-2-yl)carbamoyl)oxirane-2-carboxylate

$^{1}H$ NMR (500 MHz, $CDCl_3$)

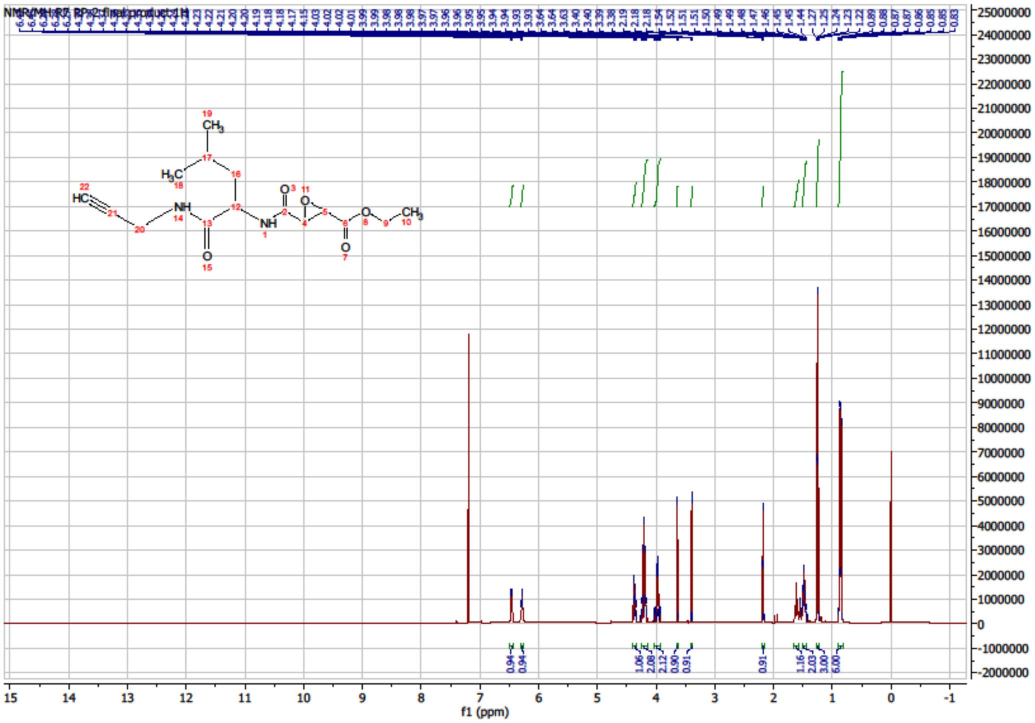

$^{13}C$ NMR {$^{1}H$} (126 MHz, $CDCl_3$)

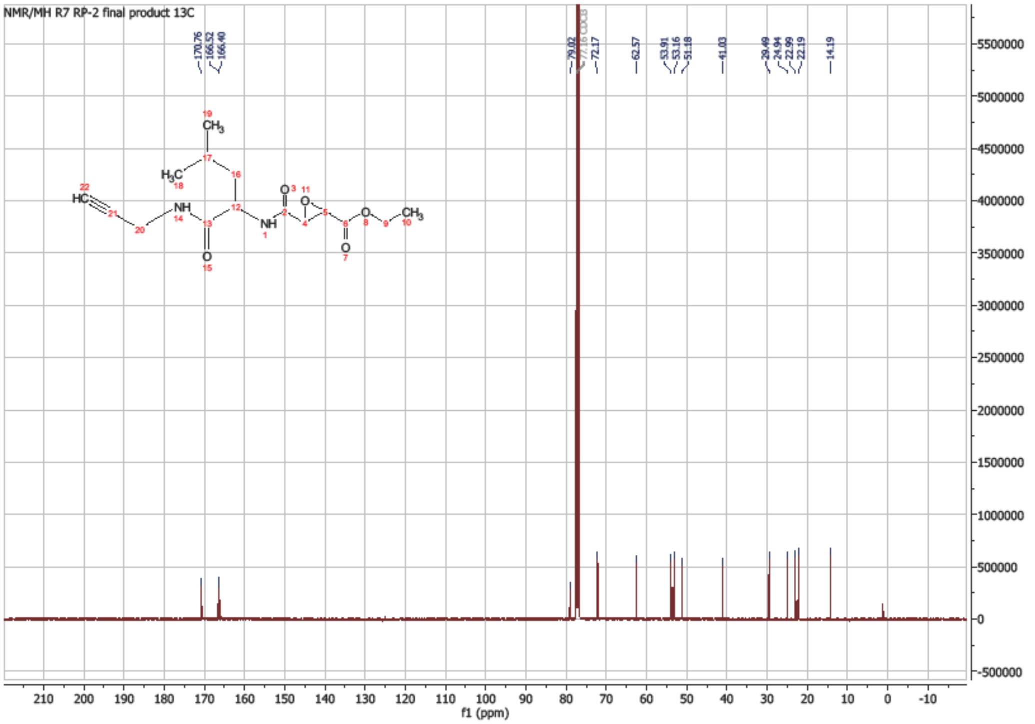

**Uncropped gels & Western Blots**

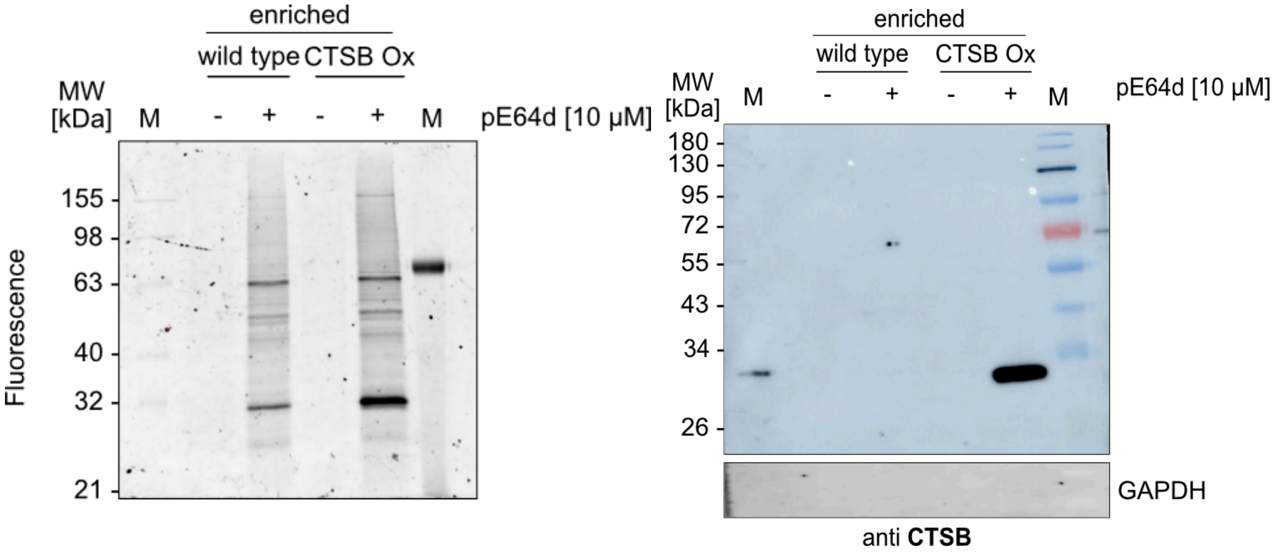

**Fig. 3a CSTB SDS PAGE & WB**

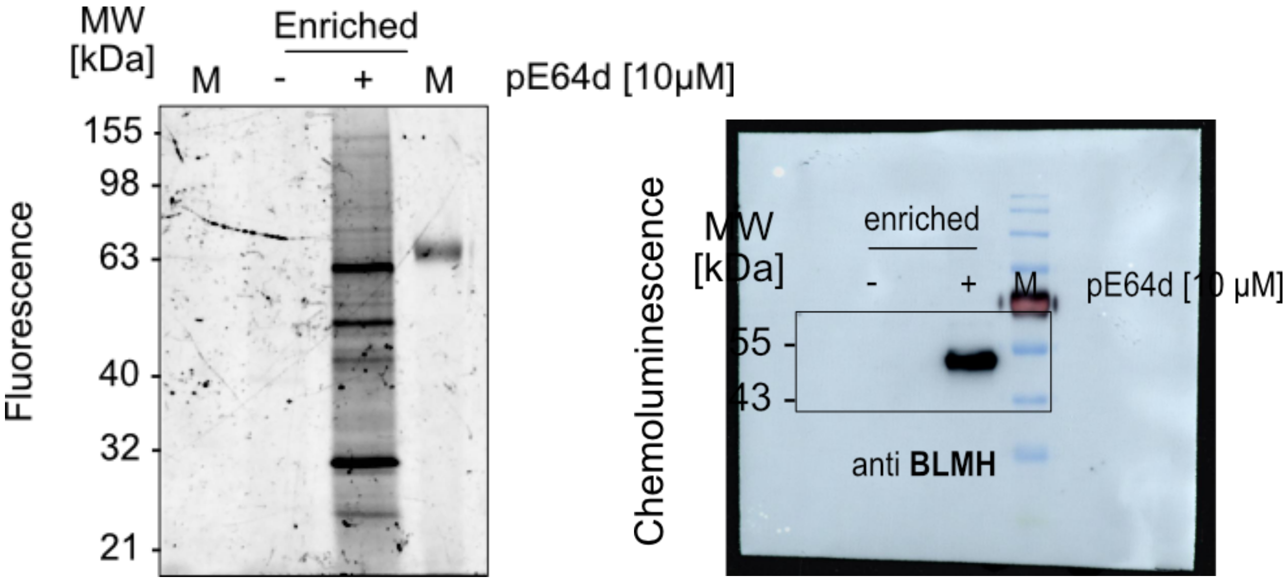

**Fig. 3b BLMH SDS PAGE & WB**

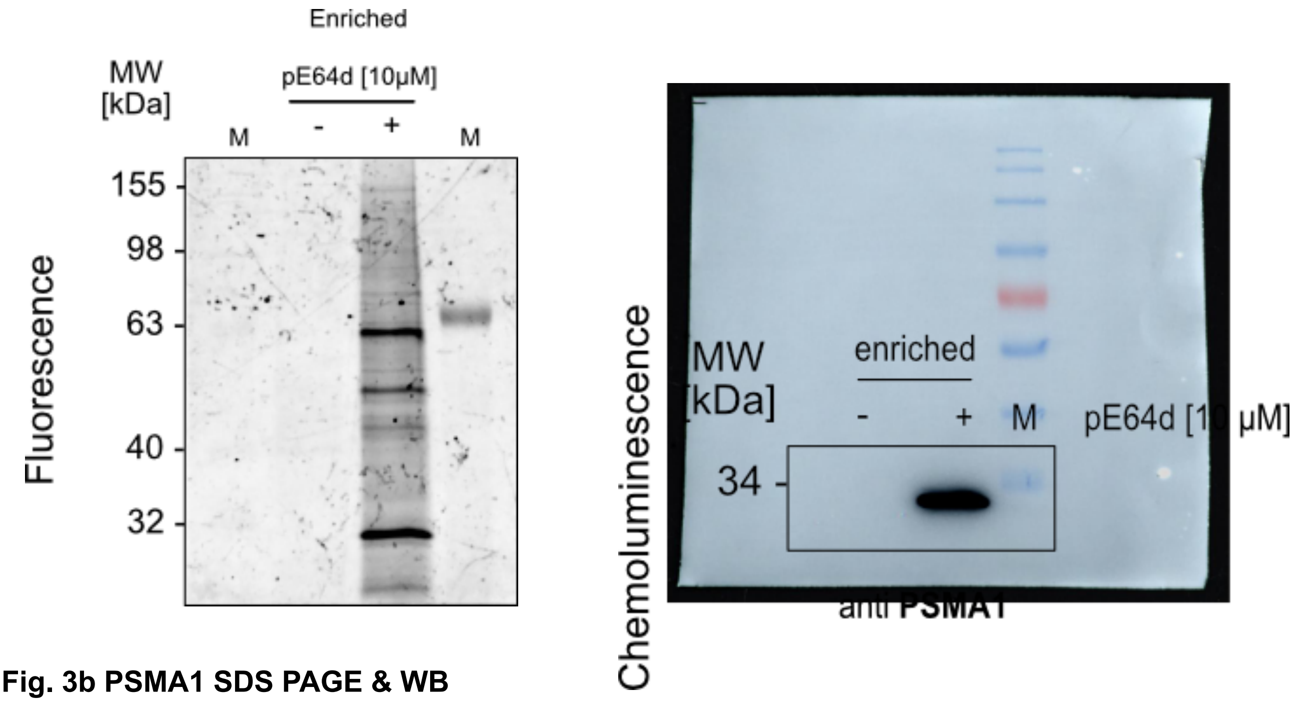

**Fig. 3b PSMA1 SDS PAGE & WB**

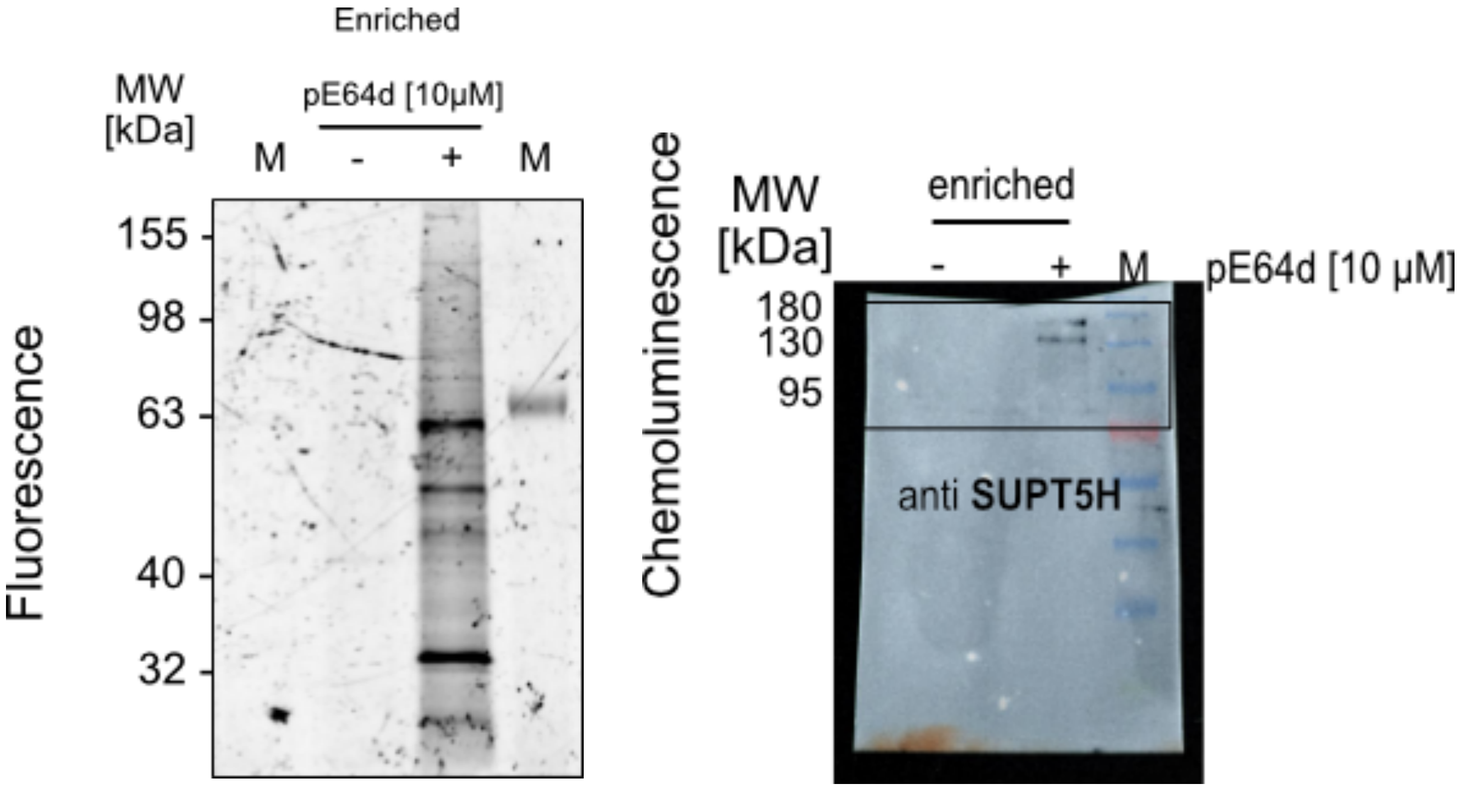

**Fig. 3b SUPT5H SDS PAGE & WB**

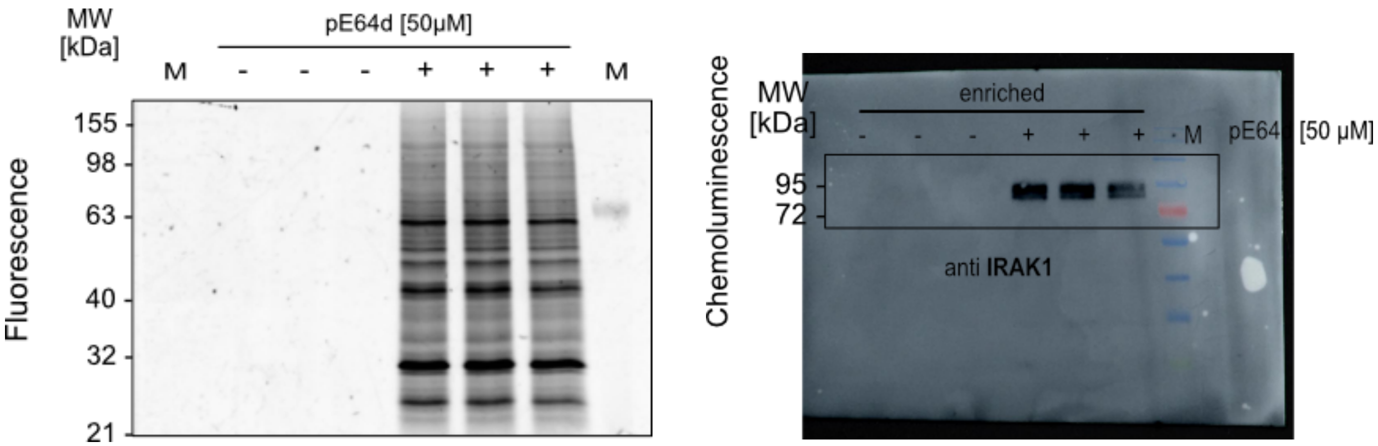

**Fig. 3b IRAK1 SDS PAGE & WB**

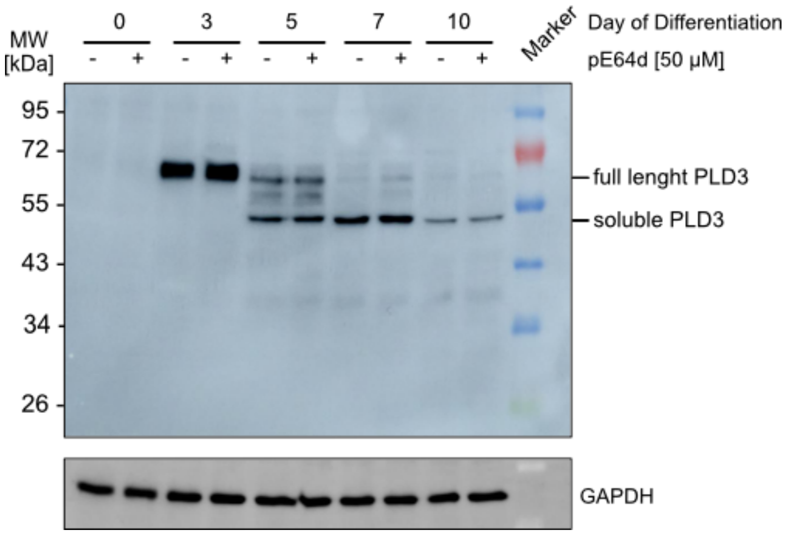

**Fig. 5b PLD3 WB**

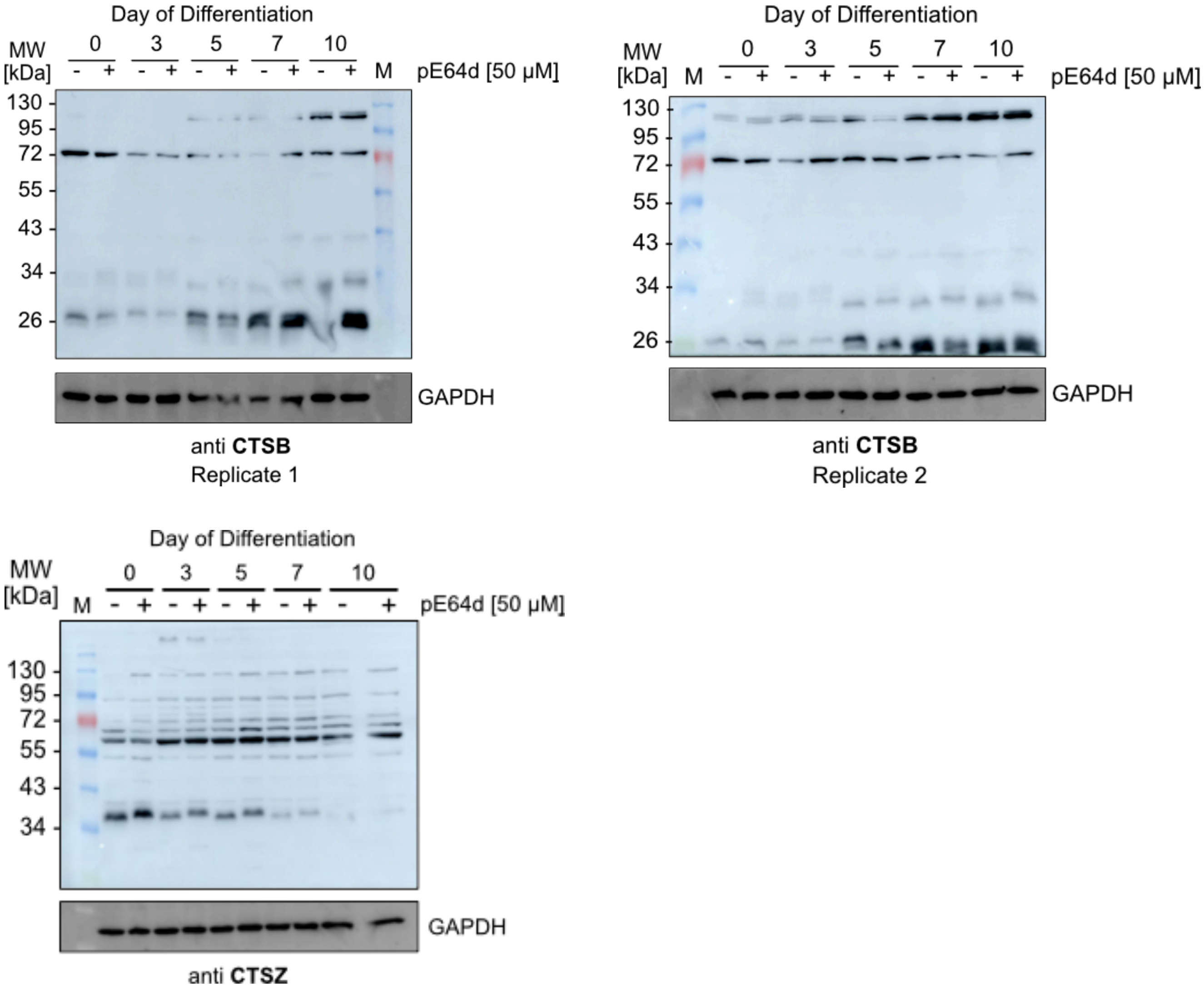

**Fig. 5b CTSB & CTSZ WB**

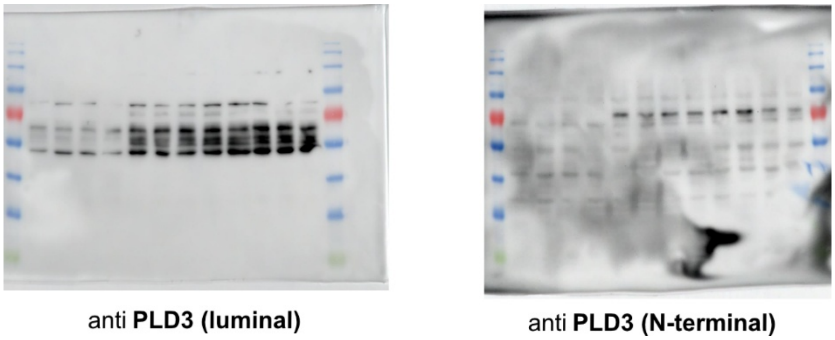

**Fig. 6a and 6b PLD3 luminal and N-terminal WB**

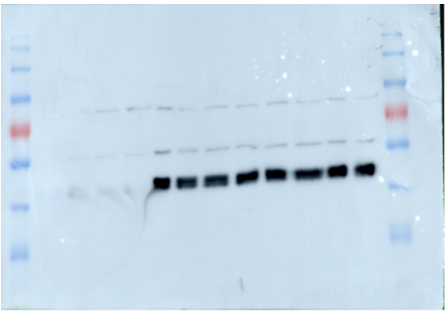

anti **PLD3 (luminal)**

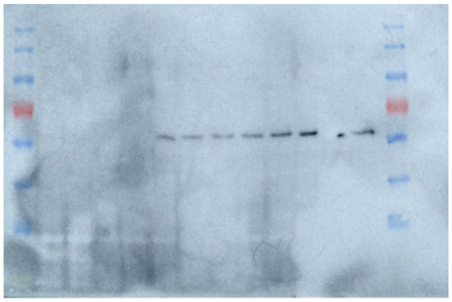

anti **PLD3 (N-terminal)**

**Fig. 6c and 6d PLD3 luminal and N-terminal WB (PNGase F treatment)**
